# Neurochemical phenotype of relaxin family peptide receptor-3 (RXFP3) lateral hypothalamus/zona incerta cells

**DOI:** 10.64898/2026.04.09.717598

**Authors:** Brandon K Richards, Jennifer L Cornish, Jee Hyun Kim, Andrew J Lawrence, Christina J Perry

## Abstract

The relaxin-3/relaxin family peptide receptor 3 (RXFP3) neuropeptidergic system is emerging as a potential target for treating various neuropsychiatric diseases, particularly those involving dysregulated stress and arousal. RXFP3 is abundantly expressed in several hypothalamic nuclei, and in the zona incerta (ZI). These regions play a central role in the regulation of stress and arousal, however the function of relaxin-3/RXFP3 within these circuits is unknown. The purpose of this study was to begin characterising this function by describing the distribution and genetic signature of neurons that express RXFP3. We used RNAscope fluorescent *in situ* hybridisation to characterise the spatial expression pattern and neurochemical phenotype of cells expressing *Rxfp3* mRNA throughout the mouse lateral hypothalamus (LH) and ZI. We found that *Rxfp3* is expressed across the rostrocaudal extent of both the LH and ZI and follows a parabolic pattern of expression, peaking in more rostral areas of each nucleus. Neurochemical phenotyping of *Rxfp3*+ cells with *Gad1*, *Slc17a6* (vGlut2), *Pvalb*, *Th*, and *Sst* showed that LH/ZI *Rxfp3*+ cells co-express each marker to varying extents, generally proportional to their overall abundance within each structure. Furthermore, LH/ZI *Rxfp3+* cells overlapped with several known populations involved in various facets of fear learning and defensive behaviour, such as the dopaminergic A13 group, somatostatin-expressing rostral ZI neurons, and glutamatergic LH neurons. The neurochemical diversity of these neurons may reflect the overall role of both the LH and ZI as global regulators of behaviour and the role of relaxin-3/RXFP3 signalling in modulating high-vigilance states.

## 1. Introduction

RXFP3 is a ligand-activated G_i/o_ protein-coupled receptor and the cognate receptor for the highly conserved neuropeptide relaxin-3 (Bathgate et al., 2006). Relaxin-3 neurons are primarily synthesised in the pontine nucleus incertus (Bathgate et al., 2002; Burazin et al., 2002; Liu et al., 2003) and project throughout the neuraxis to RXFP3-dense regions involved in arousal, stress, cognition, and appetitive motivation (Ma et al., 2007; Smith et al., 2010). We recently discovered a cell population spanning the lateral hypothalamus and zona incerta that expresses relaxin family peptide receptor-3 (LH/ZI^RXFP3^) which may regulate defensive behaviour (Richards et al., 2025). LH/ZI^RXFP3^ cells project to key regions implicated in fear learning and threat perception, including the periaqueductal gray and lateral habenula, and do so in a topographically distinct manner (Richards et al., 2026). Furthermore, following Pavlovian fear conditioning, chemogenetic activation of these cells reduced conditioned freezing behaviour and overall augmented several indices of locomotor behaviour, but induced vigorous escape-like jumping behaviour in a subset of mice (Richards et al., 2025). Given the known diversity of both nuclei, it may be that LH/ZI^RXFP3^ neurons comprise neurochemically discrete subpopulations that control distinct aspects of defensive behaviour.

The hypothalamus is a functionally diverse division of the diencephalon that is primarily involved in maintaining survival (Swanson, 1987; 2000). One essential hypothalamic role is integrating sensory information transmitted via corticostriatal pathways to coordinate appropriate defensive responses to threats (Gross & Canteras, 2012). Several medial hypothalamic nuclei, including the ventromedial hypothalamus, dorsal premammillary nucleus, and anterior hypothalamus, form highly interconnected defensive systems that respond to predators or previously fear-conditioned stimuli (Cezario et al., 2008; Dielenberg et al., 2001; Kim et al., 2017; Laing et al., 2023; Mendes-Gomes et al., 2020). The LH is more commonly associated with hunger and sleep (Braga et al., 2024; De Sousa et al., 2019; Devarakonda & Kenny, 2017; Moeyaert et al., 2018; Petzold et al., 2023; Teegala et al., 2023), but has also recently been implicated in behavioural responses to aversive stimuli (Barbano et al., 2020; Calvigioni et al., 2023; Contesse et al., 2025; Lazaridis et al., 2019; Lecca et al., 2017; Sharpe et al., 2021; Stamatakis et al., 2016).

Much less is known about the function of the ZI, an elongated structure adjacent to the hypothalamus in both human and rodent brains. Over the past decade, there has been increasing evidence that this region is central to regulating responses to aversive and threatening stimuli (Chou et al., 2018; Ho et al., 2023; Z. Li et al., 2021; Lin et al., 2023; Schroeder et al., 2023; Venkataraman et al., 2019, 2021; H. Wang et al., 2020; X. Wang, Chou, Peng, et al., 2019; Wu et al., 2023; Zhou et al., 2018). Indeed, with widespread reciprocal connectivity to sensory, emotional and motor centres in the brain, this region has been postulated to be a synaptic interface in the diencephalon (Mitrofanis, 2005; Venkataraman & Dias, 2023; X. Wang, Chou, Zhang, et al., 2019).

Both the LH and ZI contain multiple distinct cell populations with diverse neurochemical properties (Bonnavion et al., 2016; Mitrofanis, 2005). With regard to their function in threat response, cell types are typically defined and interrogated based on the expression of a single neuropeptide (e.g., hypocretin/orexin, Soya et al., 2017; W. Zhang et al., 2009), calcium-binding protein (e.g., parvalbumin; Zhou et al., 2018) receptor (e.g. RXFP3; Richards et al., 2025), or neurotransmitter (e.g. GABA; Chou et al., 2018; Venkataraman et al., 2019, 2021). However, meaningful neuronal classification is unlikely when based solely on a single cellular property (Zeng & Sanes, 2017). Indeed, subsets of GABAergic ZI neurons have been implicated in both inhibiting (Chou et al., 2018; Venkataraman et al., 2021) and promoting (Zhou et al., 2018) conditioned fear memories, suggesting that classifying ZI neurons solely by GABAergic neurotransmission may capture functionally heterogeneous populations. Therefore, further parcellation of these populations into neurochemically defined subtypes may better clarify their functional roles.

Although neurochemical phenotyping of RXFP3+ cells was initially challenging due to a lack of effective antibodies for immunohistochemical analysis, this has since been overcome by the advent of RXFP3-Cre fluorescent reporter lines (Ch’ng et al., 2019; Eraslan et al., 2024) and high-quality probes that detect *Rxfp3* mRNA *in situ*. As such, the neurochemical phenotype of RXFP3+ cells has been examined (to varying extents) using either or both methods in the bed nucleus of the stria terminalis (Ch’ng et al., 2019), hippocampus (Rytova et al., 2019), horizontal diagonal band, medial septum (Albert-Gascó et al., 2018; Haidar et al., 2019), paraventricular hypothalamus (Kania et al., 2017), retrosplenial cortex (Navarro-Sánchez et al., 2024), arcuate nucleus, dorsomedial hypothalamus, and ventral tegmental area (Voglsanger et al., 2021). These studies have consistently demonstrated that RXFP3+ cells exhibit diverse neurochemical properties; therefore, it would not be unprecedented that LH/ZI^RXFP3^ cells follow suit. Using RNAscope^®^ fluorescent *in situ* hybridisation, the current study examines the neurochemical properties of LH/ZI^RXFP3^ cells and comprehensively maps their distribution across the rostrocaudal extent of both nuclei.

## 2. Materials and Methods

### 2.1. Animals

Experiments were performed in accordance with the Prevention of Cruelty to Animals Act (2004), under the guidelines of the National Health and Medical Research Council Code of Practice for the Care and Use of Animals for Scientific Purposes (8^th^ Edition, 2013) and were approved by the Macquarie University Animal Ethics Committee (Animal Research Authority number: 2021/021). Inbred adult male RXFP3-Cre mice (*n* = 8; Ch’ng et al., 2019; Richards et al., 2025) were used for all experiments. Animals were group-housed in individually ventilated chambers and maintained on a 12-hour light-dark cycle (lights on at 6 am) in a temperature-controlled environment (21–23°C) with nesting material and *ad libitum* access to standard mouse chow and water.

### 2.2. Tissue collection and preparation

Mice were anaesthetised with an overdose of sodium pentobarbitone (80 mg/kg, i.p., Virbac, Australia) and either transcardially perfused before their brains were extracted (experiment 1), or brains were extracted immediately, frozen over dry ice, and stored at −80°C until needed (experiments 2 and 3). For experiment 1, mice were perfused transcardially using a perfusion pump at a flow rate of 7 ml/min for 2 minutes with phosphate-buffered saline (PBS; 0.1 M; pH 7.4), followed by 5 minutes of 4% w/v paraformaldehyde (PFA; Sigma-Aldrich, Australia) in 0.1 M PBS. Brains were then extracted, post-fixed in 4% PFA in 0.1 M PBS for 1 hour, washed in 0.1 M PBS for 1 hour, and immersed overnight in 20% w/v sucrose in 0.1 M PBS for cryoprotection. Finally, brains were frozen over dry ice and stored at −80°C until required. For cryosectioning, brains were placed into a Leica CM1950 Cryostat (Leica Biosystems, Germany) and allowed to equilibrate at −18°C for at least 30 minutes. Brains were embedded onto a specimen disc with Tissue-Tek Optimal Cutting Temperature compound (Sakura Finetek, USA). 17 µm (experiment 1) or 8 µm (experiments 2 and 3) coronal sections, spaced 0.16 mm apart along the rostrocaudal length of the LH/ZI were slide-mounted onto Superfrost™ Plus slides (Epredia, NH, USA). Slides were stored at −80°C until further use.

### 2.3. RNAscope^™^ fluorescent *in situ* hybridisation

RNAscope^™^ fluorescent *in situ* hybridisation was performed to determine the distribution of *Rxfp3* mRNA throughout the LH and ZI, and to examine its co-expression with *Slc17a6*, *Gad1*, *Th*, *Pvalb*, and *Sst*. Slides were removed from −80°C and underwent multiple pre-treatment steps. For 17 µm fixed-frozen sections (experiment 1), slides were rinsed in dH_2_O and baked overnight at 60°C. Sections were then treated with H_2_O_2_ (8 minutes, humid environment, room temperature (RT)), then washed twice with dH_2_O (1 minute each, RT). Slides were dried, and a hydrophobic barrier was traced around each section with an ImmEdge™ Hydrophobic Barrier PAP Pen (Vector Laboratories, Newark, USA). Slides underwent an additional bake step (60 minutes, 60°C), then were treated with a protease (Protease Plus; 20 minutes, humid environment, 40°C), and then washed twice with dH_2_O (1 minute each, RT). For 8 µm fresh-frozen sections (experiments 2 and 3), slides were fixed in 4% PFA in PBS (15 minutes, RT), washed twice in 0.1 M PBS (1 minute each, RT), and then serially dehydrated in 50% ethanol (5 minutes), 70% ethanol (5 minutes) and 100% ethanol (2x 5 minutes). Slides were dried, and a hydrophobic barrier was traced around each section. Sections underwent H_2_O_2_ treatment (10 minutes, humid environment, RT), washed twice with dH_2_O (1 minute each, RT), and then were treated with a protease (Protease III; 10 minutes, humid environment, 40°C). For both 17 µm fixed-frozen and 8 µm fresh-frozen tissue, the RNAscope^®^ Multiplex Fluorescent V2 Assay (ACDBio, USA) was performed to label *Rxfp3*, *Slc17a6*, *Gad1*, *Th*, *Pvalb*, and *Sst*. All subsequent incubation steps were performed in a humid environment at 40°C. Sections were rinsed twice with wash buffer (0.1x saline sodium citrate, 0.03% sodium dodecyl sulfate in dH_2_O) before probes for *Rxfp3* (Mm-*Rxfp3*-C2, #439381-C2; all experiments), *Slc17a6* (Mm-*slc17a6*, #319171; experiment 1), *Gad1* (Mm-*GAD1*-C3, #400951-C3; experiment 1), *Th* (Mm-*Th*, #317621; experiment 2), *Pvalb* (Mm-*Pvalb*-C3, #421931-C3; experiment 2), and *Sst* (Mm-*Sst*, #404631; experiment 3) were applied and incubated for 90 minutes. A mouse-specific positive control probe (RNAscope^®^ 3-plex Positive Control Probe-Mm, #320881) and a universal negative control probe (RNAscope^®^ 3-plex Negative Control Probe, #320871) were applied to selected sections in each experiment. Sections were then incubated in Amp1 (30 minutes), Amp2 (30 minutes), and Amp3 (15 minutes) to amplify target probes. Sections were rinsed in wash buffer after each Amp step (2x 2 minutes). Sections were then incubated in HRP C1 (15 minutes), Opal 520 (Akoya Biosciences, #FP1487001KT; 1:750; 30 minutes), and then HRP blocker (15 minutes) to develop the fluorophore signal for the probe assigned to channel 1. The same procedure was repeated for HRP C2 and HRP C3, using Opal 570 (Akoya Biosciences, #FP1488022KT; 1:750) and Opal 690 (Akoya Biosciences, #FP1497001KT; 1:750) as fluorophores, respectively. Sections were incubated with DAPI for 30 seconds, then coverslipped with Dako fluorescence mounting medium (Agilent Technologies, USA). Slides were left to dry overnight in the dark and stored at 4°C until imaging. All reagents were acquired from Advanced Cell Diagnostics, USA, unless otherwise indicated.

### 2.4. Microscopy and image acquisition

Stitched photomicrographs of sections throughout the LH/ZI were obtained using a Zeiss Axio Imager Z1 (Carl Zeiss AG, Germany) epifluorescence microscope with a NeoFluar 20x/0.5 (WD = 2.0 mm) lens. DAPI, Opal 520, Opal 570, and Opal 690 labelled excitation was provided by a Xenon HXP lamp passing through FS#49, FS#38, FS#43, and FS#50 excitation filters (Carl Zeiss AG, Germany), respectively. Images were acquired using ZEN Blue software (Carl Zeiss AG, Germany).

### 2.5. Cell quantification

Cell quantification was performed using QuPath (v0.4.2) open-source software (Bankhead et al., 2017). LH and ZI regions were manually outlined based on The Mouse Brain Atlas in Stereotaxic Coordinates (Paxinos & Franklin, 2001), and DAPI+ cells in outlined regions for all images were detected simultaneously by combining a custom script with the in-built ‘positive cell detection’ function. For the detection of *Rxfp3*, *Slc17a6*, *Gad1*, *Th*, *Pvalb*, and *Sst* mRNA and co-expression analysis, the in-built ‘object-classifier’ function in QuPath was used. Seven images from each experiment were assigned as training images for the random trees object classifier, where cells were manually assigned as ‘positive’ or ‘negative’ for the marker of interest until the machine learning algorithm produced an accurate profile of expression. To do so, a semi-quantitative method was employed, in which cells with two or more fluorescent dots within 5 µm of the DAPI-stained area were considered positive for the marker of interest (Ch’ng et al., 2019; Richards et al., 2025; Viden et al., 2022; Walker et al., 2021), with each dot denoting an individual mRNA molecule (F. Wang et al., 2012). Trained classifiers were batch-applied to all outlined regions for all images using a custom script, which classified each DAPI+ cell as being ‘positive’ or ‘negative’ for each marker. All QuPath scripts and instructions for use can be found at https://github.com/BrandonKR1.

### 2.6. Data analysis

*Rxfp3*+ counts from experiment 3 were analysed using a mixed-effects logistic regression to determine whether rostrocaudal position significantly predicted *Rxfp3*+ expression across the LH and ZI. Rostrocaudal position (in mm from Bregma) was treated as a continuous predictor, and mouse identity was included as a random intercept to account for interindividual variability in counts. Quadratic and linear model fits were compared using likelihood-ratio tests. When the quadratic model provided a significantly better fit than the linear model, the rostrocaudal position of maximal expression was estimated from the fitted coefficients.

In cases where cell composition could be stratified into more than two categories (e.g. *Rxfp3*+ cells in experiments 1 and 2) we analysed if there were rostrocaudal differences in the neurochemical identity of these cells by using multinomial logistic regressions. ZI counts were divided into four approximately equidistant rostrocaudal zones according to the Mouse Brain Atlas in Stereotaxic Coordinates (Paxinos & Franklin, 2001): the rostral zona incerta (ZIR; Bregma −0.82 to −1.34 mm), the rostral part of the ZID/ZIV (rZID/ZIV; Bregma −1.46 to −1.82 mm), the intermediate part of the ZID/ZIV (iZID/ZIV; Bregma −1.94 to −2.30 mm), and the caudal part of the ZID/ZIV + caudal zona incerta (ZIC; Bregma −2.46 to −2.92 mm). LH counts were stratified into anterior (Bregma −0.34 to −1.70 mm), tuberal (Bregma −1.82 to −2.06 mm), and mammillary (Bregma −2.18 to −2.80 mm) rostrocaudal zones according to Hahn et al. (2019). However, since the anterior division covers a large part of the hypothalamus, we reasoned that treating this area as a whole might conceal observable expression patterns within it. Therefore, the anterior LH (aLH) was further subdivided into three sectors: a rostral part (Bregma −0.34 to −0.82 mm), an intermediate part (Bregma −0.94 to −1.34 mm), and a caudal part (Bregma −1.46 to −1.70 mm). We then fitted a multinomial logistic regression model with rostrocaudal location within the LH or ZI as a categorical predictor. The overall significance of the model was evaluated using Wald’s χ^2^ test. *Post hoc* pairwise contrasts were computed on the model-estimated proportions and Sidak corrected. In cases where cell composition could be stratified into binomial categories (e.g. *Rxfp3*+ cells in experiment 3), we tested for rostrocaudal differences in the neurochemical identity of these cells using binomial logistic regressions. The overall significance of the model was evaluated with Wald’s χ^2^ test, and *post hoc* pairwise contrasts were computed on the model-estimated proportions. *p* < .05 was considered statistically significant.

To generate schematics illustrating the spatial distribution of *Rxfp3* mRNA and its co-expression with select neurochemical markers, cell classification masks were exported from QuPath as .svg files and transposed onto the corresponding Mouse Brain Atlas in Stereotaxic Coordinates plate (Paxinos & Franklin, 2001) using Adobe Illustrator (Adobe, San Jose, USA) software. Regression analyses were run, and regression graphs were made in STATA. All other graphs were created in PRISM 7 (GraphPad Software, San Diego, USA). Adobe Illustrator was used to construct figures.

## 3. Results

### 3.1. Distribution of *Rxfp3* across the LH and ZI

Overall, 18.0% (± 2.1%) of DAPI+ cells in the ZI and 15.9% (± 1.9%) in the LH were *Rxfp3*+ (Figure 1A, B). A mixed-effects logistic regression showed that rostrocaudal location significantly predicted *Rxfp3* expression in both the LH and ZI. A quadratic model fit significantly better than a linear model in both nuclei (ZI: likelihood-ratio test, χ^2^_(1)_ = 151.0, *p* < .001; LH: likelihood-ratio test, χ^2^_(1)_ = 696.56, *p* < .001). In the ZI, the fitted curve peaked at −1.41 mm from Bregma in the rZID/ZIV (95% CI: −1.51 to −1.32 mm; Figure 1C), whereas in the LH, the curve peaked at −1.47 mm from Bregma, bordering the intermediate and caudal aLH (95% CI: −1.50 to −1.45 mm; Figure 1D). These results indicate that *Rxfp3* expression in the LH and ZI follows a parabolic pattern, peaking in the more rostral regions of each nucleus. We then mapped *Rxfp3*+ cells across the LH and ZI to examine spatial expression patterns. *Rxfp3*+ cells were evenly distributed throughout the ZIR (Figure 1E - G, M), whereas in the rZID/ZIV, they primarily occupied the medial ZIV, especially surrounding the mammillothalamic tract (Figure 1H, N). Throughout the rest of the ZI, *Rxfp3* expression was strongly biased toward the ZIV rather than the ZID (Figure 1I - L, O). Clusters of *Rxfp3*+ cells were observed in the lateral ZIV, contiguous with *Rxfp3*+ cells in the subgeniculate and ventrolateral geniculate nuclei of the thalamus (Figure 1O). In the LH, although a cluster of *Rxfp3*+ cells was observed surrounding the fornix in the tuberal region (Figure 1I), *Rxfp3*+ cells were evenly distributed across the LH.

**Figure 1.**
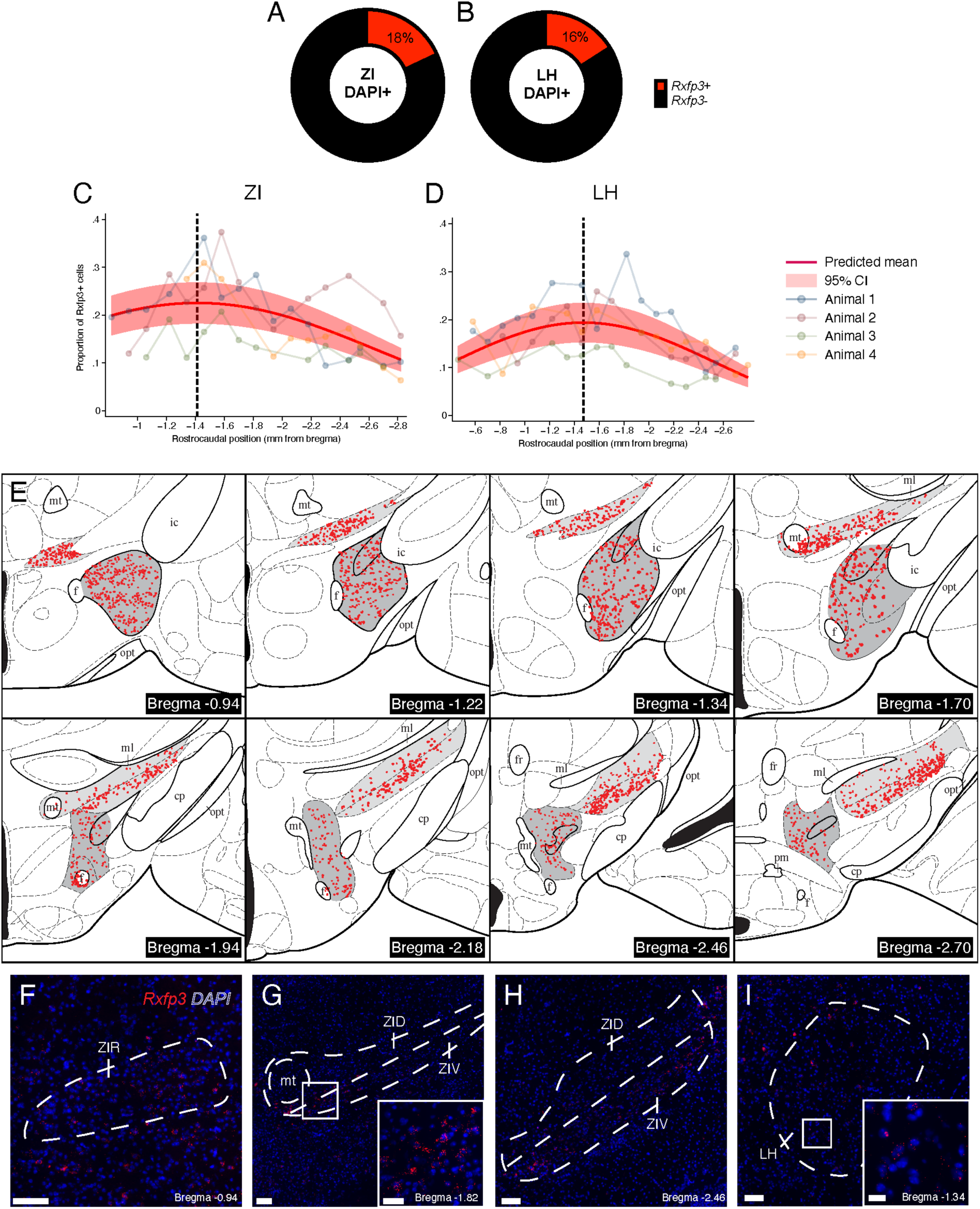
Distribution of *Rxfp3* mRNA across the LH and ZI. **(A-B)** Donut chart quantification of the mean proportion of DAPI+ cells expressing *Rxfp3* (*Rxfp3*+, red) or not (*Rxfp3*-, black) in the ZI (A) and LH (B) **(C-D)** Graphs showing the raw proportions of DAPI+ cells positive for *Rxfp3* mRNA expression along the rostrocaudal axis of the ZI (C) and LH (D; transparent dots and lines), overlaid with the fitted predicted means and 95% confidence intervals from the mixed model logistic regressions (solid red lines and shading). Dotted black lines indicate the predicted peak of expression. **(E)** Schematics of *Rxfp3* expression throughout the LH (dark grey) and ZI (light grey) mapped onto plates from the Mouse Brain Atlas in Stereotaxic Coordinates (Paxinos & Franklin, 2001) from a representative RXFP3-Cre mouse. **(F-I)** Representative photomicrographs of RNAscope^®^ fluorescent *in situ* hybridisation for *Rxfp3* mRNA in the rostral zona incerta (F), intermediate zona incerta (G), caudal part of the intermediate zona incerta (G), and the LH (H). Note the dense cluster of *Rxfp3*+ cells surrounding the mammillothalamic tract in F, shown in high-power as an inset image in the bottom right corner. The inset image in I shows a typical *Rxfp3*+ expression pattern throughout the LH. Inset image scale bar in F = 50 µm. Inset image scale bar in H = 25 µm. All other scale bars = 100 µm. Abbreviations: cp, cerebral peduncle; f, fornix; fr, fasciculus retroflexus; ic, internal capsule; LH, lateral hypothalamus; ml, medial lemniscus; mt, mammillothalamic tract; opt, optic tract; pm, principal mammillary tract; ZIR, rostral zona incerta; ZID, dorsal zona incerta; ZIV, ventral zona incerta.

### 3.2. Neurochemical phenotyping of *Rxfp3*+ cells across the LH and ZI

#### 3.2.1. Experiment 1: *Slc17a6* and *Gad1*

First, we were interested in the fast-acting, amino-acid neurotransmitter phenotype of LH/ZI^RXFP3^ cells. Therefore, we examined the co-expression of *Rxfp3* with *Slc17a6* (vGlut2, a marker of glutamatergic neurotransmission) and *Gad1* (GAD1, a marker of GABAergic neurotransmission). Overall, ZI *Rxfp3*+ cells were primarily *Gad1*+ (82.1% ± 4.0%), and were rarely *Slc17a6*+ (1.4% ± 0.7%; Figure 2A, M). However, only 25.9% (± 1.0%) of ZI *Gad1*+ cells were *Rxfp3*+ (Figure 2H), indicating that *Rxfp3*+/*Gad1*+ cells were only a subset of ZI *Gad1*+ cells. Despite the low overall number of ZI *Rxfp3*+/*Slc17a6*+ cells, 34.6% (± 5.5%) of ZI *Slc17a6*+ cells were *Rxfp3*+ (Figure 2E). Overall, this data suggests that *Rxfp3* was not selectively expressed on either ZI *Gad1*+ or *Slc17a6*+ neurons. However, as most of the ZI was *Gad1*+, *Rxfp3*+/*Gad1*+ cells were more abundant than *Rxfp3*+/*Slc17a6*+ cells. In the LH, 26.4% (± 2.9%) of *Rxfp3*+ cells were *Gad1*+ and 35.2% (± 5.8%) were *Slc17a6*+ (Figure 2C). 24.4% (± 2.8%) of LH *Gad1*+ cells co-expressed *Rxfp3* (Figure 2L), while 38.1% (± 3.0%) of LH *Slc17a6*+ cells co-expressed *Rxfp3* (Figure 2I). Taken together, LH *Rxfp3*+ cells expressed *Gad1*+ and *Slc17a6*+ in relatively equal proportions and were subsets of LH *Gad1*+ and LH *Slc17a6*+ cells.

**Figure 2.**
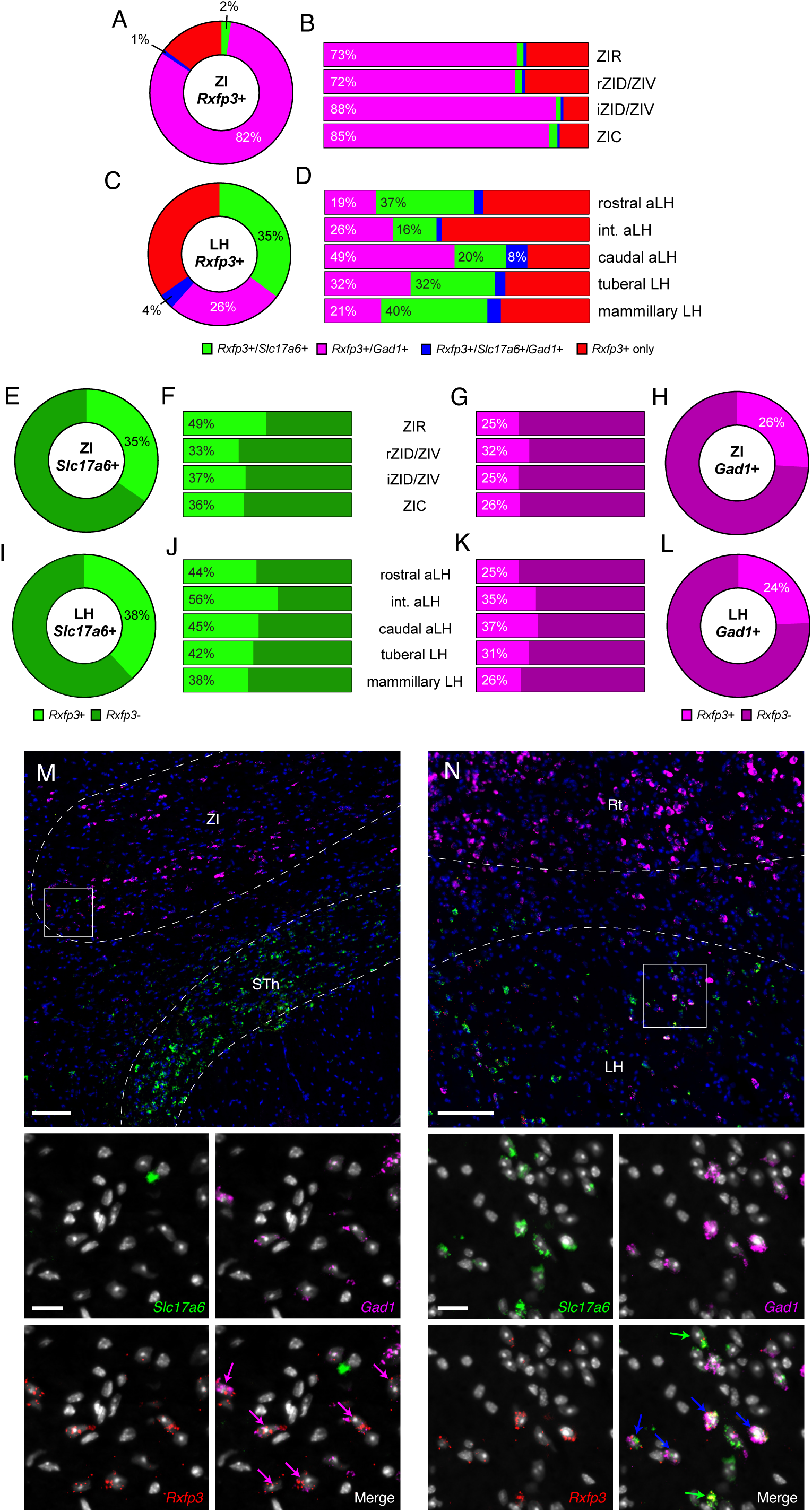
Co-localisation of *Rxfp3* with *Slc17a6* and *Gad1* throughout the ZI and LH. **(A, C)** Donut chart quantification of the mean proportion of *Rxfp3*+ cells expressing *Slc17a6* (*Rxfp3*+/*Slc17a6*+, green), *Gad1* (*Rxfp3*+/*Gad1*+, magenta) both (*Rxfp3*+/*Slc17a6*+/*Gad1*+, blue), or no co-expression (*Rxfp3*+ only, red) in the ZI (A) and LH (C). **(B, D)** Distribution plots showing the mean proportion of *Rxfp3*+ cells expressing *Slc17a6*, *Gad1*, both, or neither, but split across rostrocaudal location for the ZI (B) and LH (D). Reported means are adjusted values based on multinomial logistic regression. **(E, I)** Donut chart quantification of the mean proportion of *Slc17a6*+ cells expressing *Rxfp3* (*Rxfp3*+, light green) or not (*Rxfp3*-, dark green) for both the ZI (E) and LH (I). **(F, J)** Distribution plots showing the mean proportion of *Slc17a6*+ cells expressing *Rxfp3* or not, but split across rostrocaudal location for the ZI (F) and the LH (J). Reported means are adjusted values based on binomial logistic regression. **(G, K)** Distribution plots showing the mean proportion of *Gad1*+ cells expressing *Rxfp3* or not, but split across rostrocaudal location for the ZI (G) and the LH (K). Reported means are adjusted values based on binomial logistic regression. (**H, L)** Donut chart quantification of the mean proportion of *Gad1*+ cells expressing *Rxfp3* (*Rxfp3*+, magenta) or not (*Rxfp3*-, purple) for both the ZI (H) and LH (L). **(M)** Representative fluorescent photomicrograph of cells expressing *Rxfp3*, *Slc17a6*, and *Gad1* relative to DAPI-stained nuclei in the ZI. Zoomed in images show the inset box in the top panel. *Rxfp3*+/*Gad1*+ cells are indicated by magenta arrows. Note the predominant *Gad1*+ cell population compared to the *Slc17a6*+ population, which is more evident in the adjacent STh. **(N)** Representative fluorescent photomicrograph of cells expressing *Rxfp3*, *Slc17a6* and *Gad1* relative to DAPI-stained nuclei in the LH. Zoomed in images show the inset box in the top panel, demonstrating a cluster of triple-labelled *Rxfp3*+/*Slc17a6*+/*Gad1*+ cells (blue arrows). *Rxfp3*+/*Slc17a6*+ cells are indicated by green arrows. Main image scale bars = 100 µm. Inset image scale bars = 20 µm. All *n*s = 4 mice.

In the ZI, a multinomial logistic regression revealed a significant overall effect of rostrocaudal location on *Rxfp3*+ composition (χ^2^_(3)_ = 29.33, *p* < .001; Figure 2B). *Post-hoc* pairwise comparisons revealed that the proportion of *Rxfp3*+/*Gad1*+ cells was significantly lower in the rZID/ZIV compared to the iZID/ZIV (*p* < .001) and ZIC (*p* = .016). No pairwise comparisons involving *Rxfp3*+*/Slc17a6*+ or *Rxfp3*+*/Gad1*+*/Slc17a6*+ were statistically significant (all *p*s > .05). Overall, *Rxfp3*+ cells expressed more *Gad1*+ in the caudal half of the ZI compared to the rostral half, but the proportions of *Slc17a6*+ and *Gad1*+*/Slc17a6*+ co-expressing cells remained stable. In the LH, there was also a significant overall effect of rostrocaudal location on *Rxfp3*+ composition (χ^2^_(3)_ = 23.51, *p* < .001; Figure 2D). *Post-hoc* pairwise comparisons demonstrated that *Rxfp3/Gad1* expression was significantly higher in the caudal aLH compared with the rostral aLH, mammillary LH, and tuberal LH (all *p*s < .001). Additionally, *Rxfp3/Slc17a6* expression was significantly higher in the mammillary LH compared to the intermediate aLH (*p* = .003). Notably, the proportion of *Rxfp3*+/*Gad1*+/*Slc17a6*+ cells was significantly higher in the caudal aLH compared to the intermediate aLH (*p* < .001) and the rostral aLH (*p* = .004).

Next, we used binomial logistic regressions to test whether the proportion of *Gad1*+ and *Slc17a6*+ cells expressing *Rxfp3* varied along the rostrocaudal axis of the LH and ZI. For ZI *Gad1*+ cells, there was a significant overall effect of rostrocaudal location on *Rxfp3* co-expression (^2^_(3)_ = 431.07, *p* < .001). *Post hoc* pairwise comparisons demonstrated that the proportion of *Gad1*+/*Rxfp3*+ cells was higher in the rZID/ZIV than in the ZIR (*p* < .001) and the iZID/ZIV (*p* = .010). For ZI *Slc17a6*+ cells, there was also a significant overall effect (χ^2^_(3)_ = 12.53, *p* = .0058), but *post hoc* pairwise comparisons did not reveal any statistically significant differences (all *p*s > .05). For LH *Gad1*+ cells, there was a significant overall effect of rostrocaudal location on *Rxfp3* co-expression (χ^2^_(3)_ = 14.21, *p* = .0026; Figure 2K). *Post-hoc* pairwise comparisons indicated that the proportion of *Gad1*+/*Rxfp3*+ cells was higher in the caudal aLH than in the rostral aLH (*p* < .001), and higher in the intermediate aLH than in both the mammillary LH (*p* = .007) and the rostral aLH (*p* = .01). For LH *Slc17a6*+ cells, there was no significant effect of rostrocaudal location on *Rxfp3* co-expression (χ^2^_(3)_ = 2.85, *p* = .4146; Figure 2J).

We then examined the spatial distribution of *Rxfp3*+/*Slc17a6*+, *Rxfp3*+/*Gad1*+, and *Rxfp3*+/*Slc17a6*+/*Gad1*+ cells across the LH and ZI. In the ZI, *Rxfp3*+/*Gad1*+ cells densely occupied the ZIR (Figure 3A, B) and the medial ZIV adjacent to the mammillothalamic tract (Figure 3C, D). *Rxfp3*+/*Slc17a6*+ cells were sparsely spread throughout the ZI; however, some selectively populated the medial ZID (Figure 3D) and more caudal areas (Figure 3F). In the LH, *Rxfp3*+/*Slc17a6*+ cells outnumbered *Rxfp3*+/*Gad1*+ cells at its rostral pole (Figure 3G). However, *Rxfp3*+/*Slc17a6*+ and *Rxfp3*+/*Gad1*+ cells were generally intermingled in roughly equal proportions throughout the rest of the rostral aLH and the entire intermediate aLH (Figure 3H – J). In the caudal part of the aLH, *Rxfp3*+/*Slc17a6*+ and *Rxfp3*+/*Gad1*+ cells formed two partially overlapping clusters. The *Rxfp3*+/*Slc17a6*+ cluster occupied the ventrolateral LH adjacent to the optic tract, while the *Rxfp3*+/*Gad1*+ cluster occupied the medial perifornical area and extended into the dorsomedial LH, overlapping the nigrostriatal bundle (Figure 3K, L). At the boundary of these populations, a significant number of *Rxfp3*+/*Slc17a6*+/*Gad1*+ cells were observed (Figure 3K, L, Figure 2N). Clustering of *Rxfp3*+/*Slc17a6*+ and *Rxfp3*/*Gad1*+ populations was absent at the tuberal level and resembled the pattern observed in the rostral aLH (Figure 3M). At the mammillary level, *Rxfp3*+/*Slc17a6*+ cells predominated in a small area immediately dorsal to the fornix, whereas *Rxfp3*+/*Gad1*+ cells were generally absent from this region (Figure 3N).

**Figure 3.**
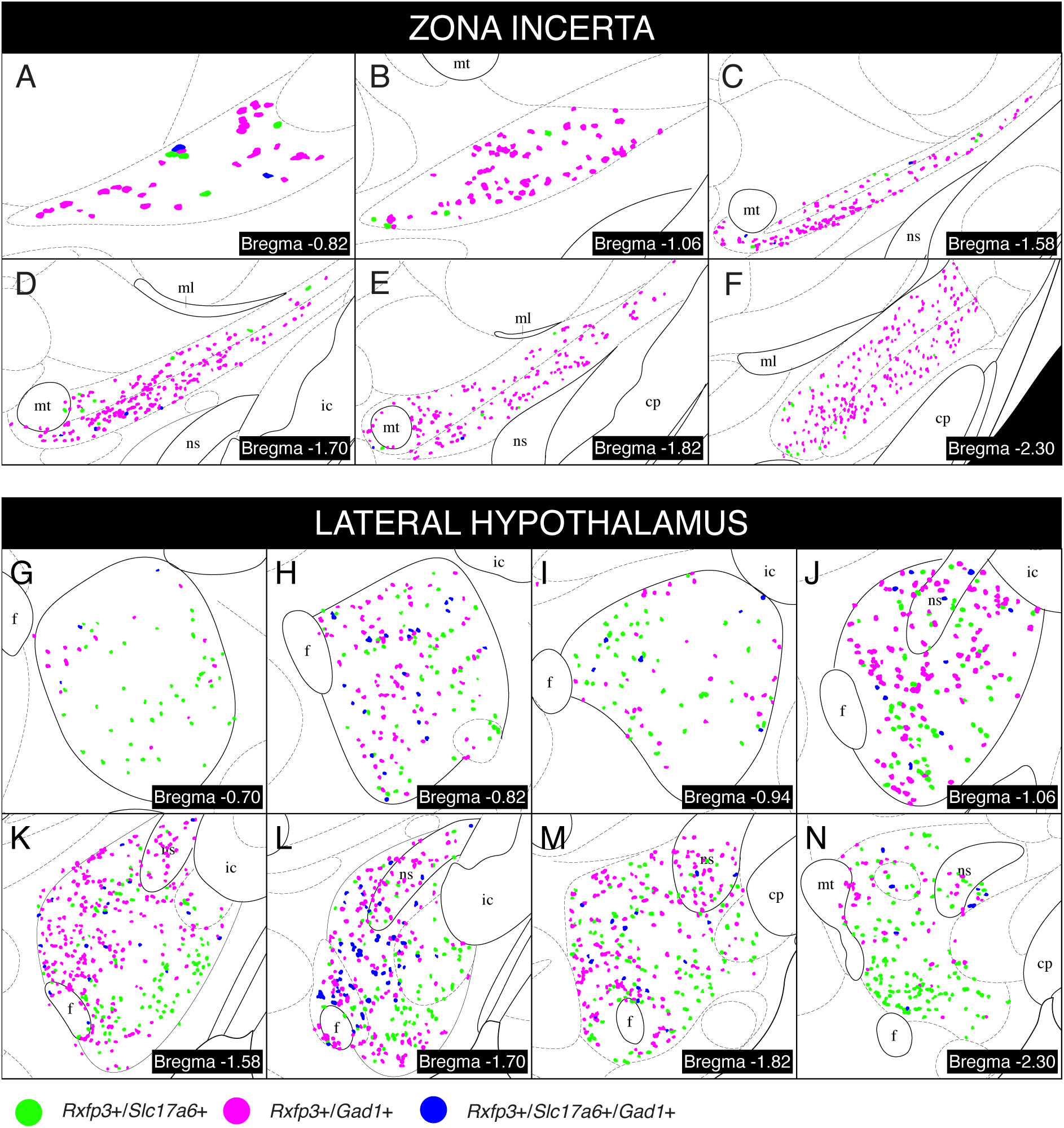
Schematics of *Rxfp3* co-localisation with *Slc17a6* and *Gad1* throughout the ZI and LH. *Rxfp3*+ cells that co-expressed *Slc17a6* (green dots), *Gad1* (magenta dots), or both *Slc17a6* and *Gad1* (blue dots) were mapped onto plates within the ZI (A – F) and LH (G – N) from the Mouse Brain Atlas in Stereotaxic Coordinates (Paxinos & Franklin, 2001) from a representative RXFP3-Cre mouse. Abbreviations: cp, cerebral peduncle; f, fornix; ic, internal capsule; ml, medial lemniscus; mt, mammillothalamic tract; ns, nigrostriatal bundle.

#### 3.2.2. Experiment 2: Tyrosine hydroxylase and parvalbumin

Recently, populations of tyrosine hydroxylase-expressing (TH+) neurons in the ZI and parvalbumin-expressing (PV+) neurons in the LH and ZI have separately been linked to the regulation of fear learning, defensive behaviours, and aversive processing (Siemian et al., 2019; Venkataraman et al., 2021; Zhou et al., 2018). Therefore, we examined whether LH/ZI^RXFP3^ cells overlapped with these populations by assessing the co-expression of *Rxfp3* with *Th* and *Pvalb* mRNA. We first examined a known cluster of TH+ cells in the rostromedial ZI, the A13 dopaminergic group (Figure 4). 55.0% (± 7.5%) of A13 *Rxfp3*+ cells were *Th*+ (Figure 4A), and 31.5% (± 8.2%) of A13 *Th*+ cells were *Rxfp3+* (Figure 4B). This indicates that a large subset of A13 RXFP3-expressing cells is dopaminergic, but many dopaminergic A13 cells do not express RXFP3. Across the entire ZI, only 7.3% (± 1.3%) of *Rxfp3*+ cells were *Th*+ (Figure 5A). However, 33.6% (± 4.6%) of *Th*+ cells were *Rxfp3*+, similar to the A13 (Figure 4B, 5E). 45.3% (± 5.8%) of ZI *Rxfp3*+ cells were *Pvalb*+ (Figure 5A), and 30.1% (± 4.1%) of *Pvalb*+ cells were *Rxfp3*+ (Figure 5H), suggesting that a substantial subset of ZI^RXFP3^ cells express PV and vice-versa. Only 0.7% (± 0.2%) of *Rxfp3*+ cells were both *Th*+ and *Pvalb*+, indicating that ZI *Rxfp3*+/*Th*+ and *Rxfp3*+/*Pvalb*+ cells are separate populations (Figure 5A). In the LH, 17.9% (± 2.1%) of *Rxfp3*+ cells were *Th*+ and 12.6% (± 5.0%) were *Pvalb*+ (Figure 5C), indicating that LH^RXFP3^ cells consist of TH+ and PV+ populations in relatively equal proportions. 39.6% (± 4.4%) of LH *Th*+ cells were *Rxfp3*+ (Figure 5I), while 19.0% (± 3.6%) of LH *Pvalb*+ cells were *Rxfp3*+ (Figure 5L). This indicates that TH+/RXFP3+ cells constitute a large subset of LH TH+ cells, whereas PV+/RXFP3+ cells comprise a smaller subset of LH PV+ cells.

**Figure 4.**
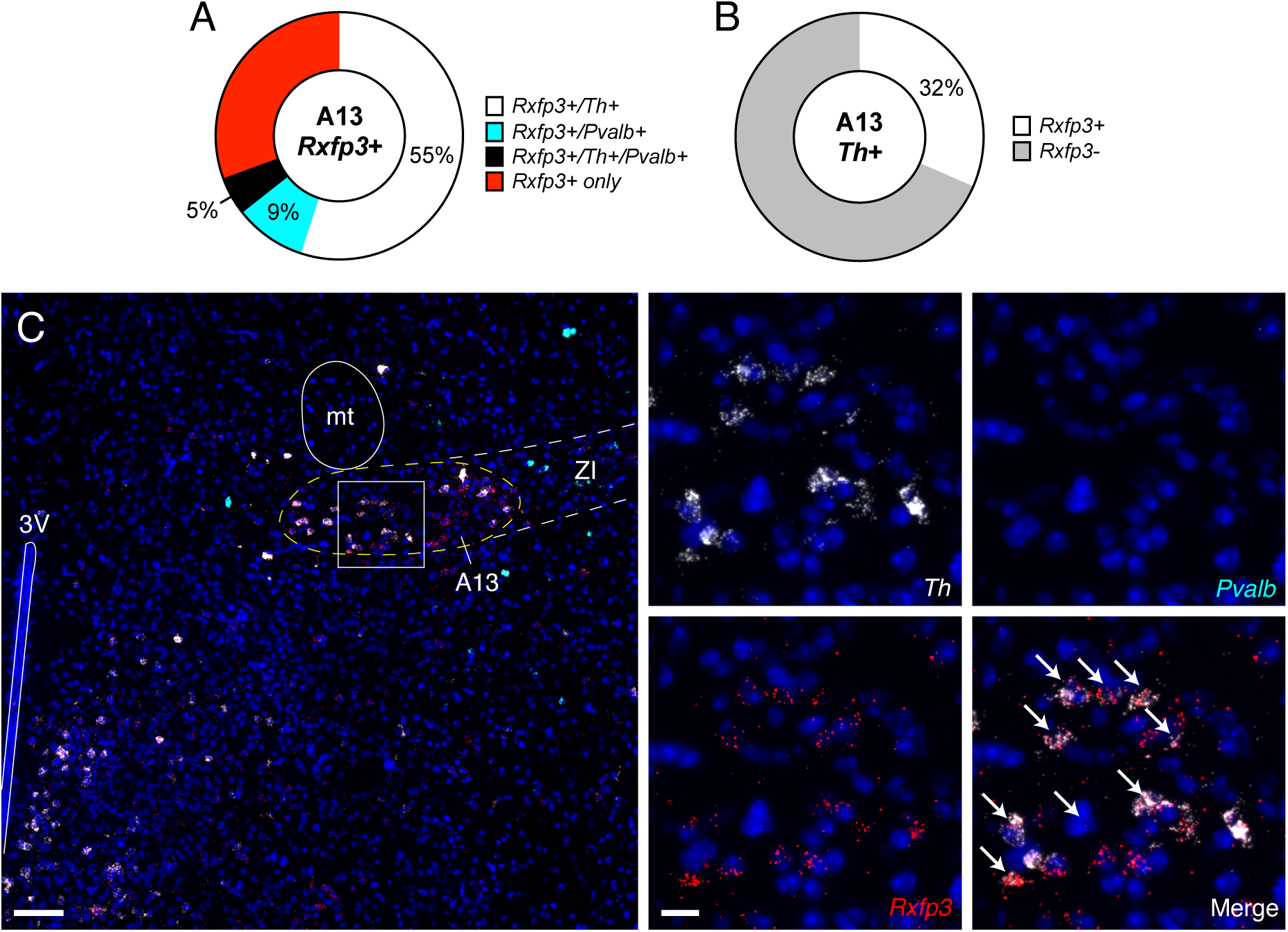
Co-localisation of *Rxfp3* with *Th* and *Pvalb* in the A13. **(A)** Donut chart quantification of the mean proportion of *Rxfp3*+ cells expressing *Th* (*Rxfp3*+/*Th*+, white), *Pvalb* (*Rxfp3*+/*Pvalb*+, cyan) both (*Rxfp3*+/*Th*+/*Pvalb*+, black), or no co-expression (*Rxfp3*+ only, red) in the A13. **(B)** Donut chart quantification of the mean proportion of *Th*+ cells expressing *Rxfp3* (*Rxfp3*+, white) or not (*Rxfp3*-, grey) for the A13. **(C)** Representative fluorescent photomicrograph of cells expressing *Rxfp3*, *Th*, and *Pvalb* mRNA relative to DAPI-stained nuclei in the A13. Zoomed in images show the inset box in the left panel. *Rxfp3*+/*Th*+ cells are indicated by white arrows. Main image scale bar = 100 µm. Inset image scale bar = 20 µm. All *n*s = 4 mice.

**Figure 5.**
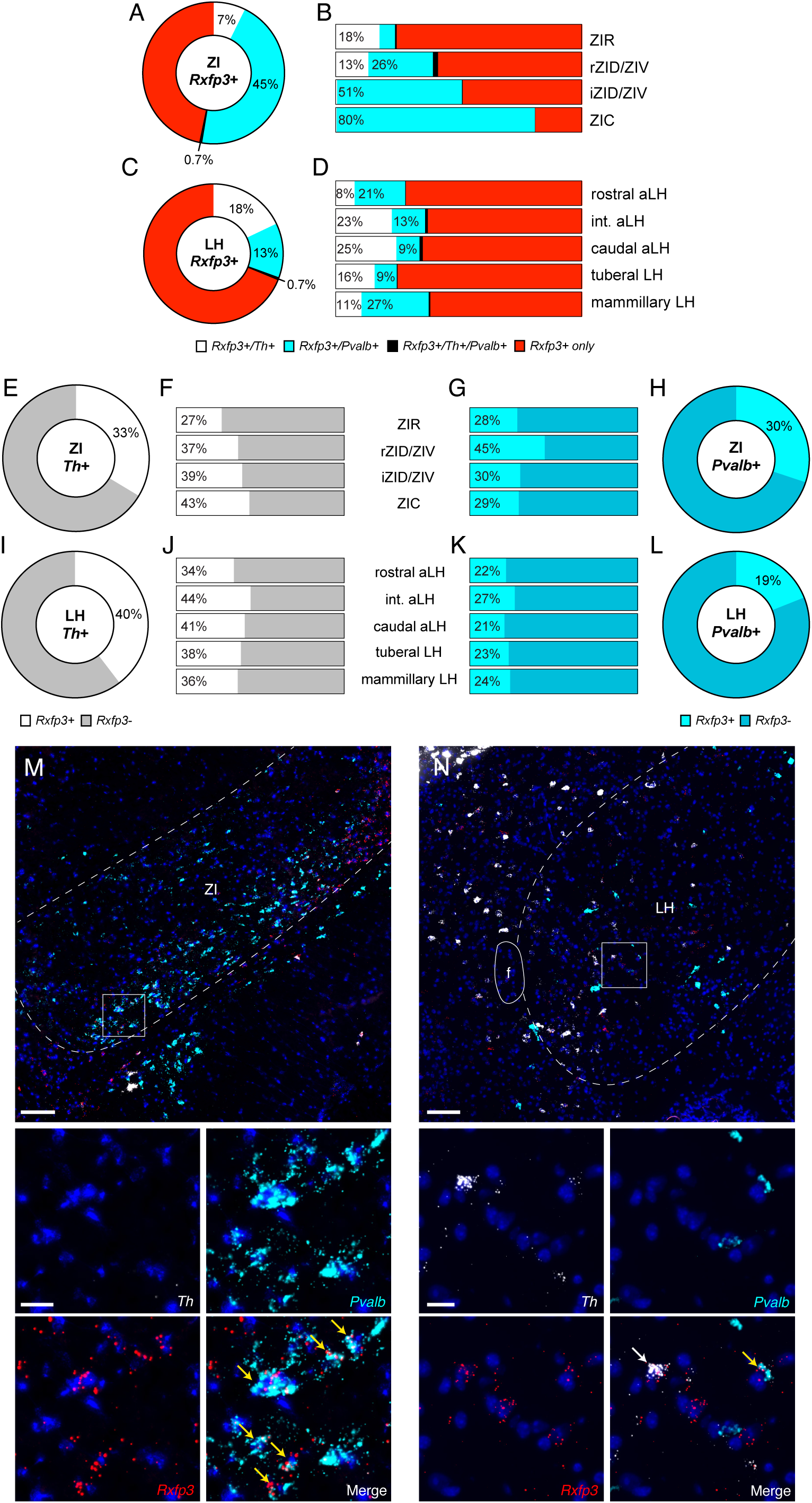
Co-localisation of *Rxfp3* with *Th* and *Pvalb* throughout the ZI and LH. **(A, C)** Donut chart quantification of the mean proportion of *Rxfp3*+ cells expressing *Th* (*Rxfp3*+/*Th*+, white), *Pvalb* (*Rxfp3*+/*Pvalb*+, cyan) both (*Rxfp3*+/*Th*+/*Pvalb*+, black), or no co-expression (*Rxfp3*+ only, red) in the ZI (A) and LH (C). **(B, D)** Distribution plots showing the mean proportion of *Rxfp3*+ cells expressing *Th*, *Pvalb*, both, or neither, but split across rostrocaudal location for the ZI (B) and LH (D). Reported means are adjusted values based on multinomial logistic regression. **(E, I)** Donut chart quantification of the mean proportion of *Th*+ cells expressing *Rxfp3* (*Rxfp3*+, white) or not (*Rxfp3*-, grey) for both the ZI (E) and LH (I). **(F, J)** Distribution plots showing the mean proportion of *Th*+ cells expressing *Rxfp3* or not, but split across rostrocaudal location for the ZI (F) and the LH (J). Reported means are adjusted values based on binomial logistic regression. **(G, K)** Distribution plots showing the mean proportion of *Pvalb*+ cells expressing *Rxfp3* or not, but split across rostrocaudal location for the ZI (G) and the LH (K). Reported means are adjusted values based on binomial logistic regression. (**H, L)** Donut chart quantification of the mean proportion of *Pvalb*+ cells expressing *Rxfp3* (*Rxfp3*+, cyan) or not (*Rxfp3*-, blue) for both the ZI (H) and LH (L). **(M)** Representative fluorescent photomicrograph of cells expressing *Rxfp3*, *Th*, and *Pvalb* mRNA relative to DAPI-stained nuclei in the ZI. Zoomed in images show the inset box in the top panel. *Rxfp3*+/*Pvalb*+ cells are indicated by yellow arrows. Note the predominant *Rxfp3*+/*Pvalb*+ cell population compared to the *Rxfp3*+/*Th*+ population. **(N)** Representative fluorescent photomicrograph of cells expressing *Rxfp3*, *Th* and *Pvalb* mRNA relative to DAPI-stained nuclei in the LH. Zoomed in images show the inset box in the top panel. An Rxfp3+/Pvalb+ cell is indicate by a yellow arrow; an Rxfp3+/Th+ cell is indicated by a white arrow. Main image scale bars = 100 µm. Inset image scale bars = 20 µm. All *n*s = 4 mice.

In the ZI, a multinomial logistic regression revealed a significant overall effect of rostrocaudal location on *Rxfp3*+ composition (χ^2^_(3)_ = 407.14, *p* < .001; Figure 5B). *Post hoc* pairwise comparisons showed that the proportion of *Rxfp3*+/*Th*+ cells was significantly higher in the ZIR and rZID/ZIV than in the iZID/ZIV and ZIC (all *p*s < .001). Additionally, all pairwise comparisons for *Rxfp3*+/*Pvalb*+ cells were statistically significant (all ps < .05). Further analysis indicated that *Rxfp3*+/*Pvalb*+ proportions increased with rostrocaudal location (Figure 5B). In the LH, there was also a significant overall effect of rostrocaudal location on *Rxfp3*+ composition (χ^2^_(3)_ = 11.04, *p* = .0115; Figure 5D). *Post hoc* pairwise comparisons showed that the proportion of *Rxfp3*+/*Th*+ cells was significantly higher in the caudal aLH and intermediate aLH than in the mammillary LH and rostral aLH (caudal aLH vs mammillary LH, caudal aLH vs rostral aLH, intermediate aLH vs rostral aLH, *p*s < .001; intermediate aLH vs mammillary LH, *p* = .008). Additionally, the proportion of *Rxfp3*+/*Pvalb*+ cells was significantly higher in the mammillary LH than in the caudal aLH (*p* < .001) and the tuberal LH (*p* = .002; Figure 5D).

Next, we used binomial logistic regressions to test whether the proportion of *Th*+ and *Pvalb*+ cells expressing *Rxfp3* varied along the rostrocaudal axis of the LH and ZI. For ZI *Th*+, LH *Th*+, and LH *Pvalb*+ cells, there was a significant overall effect of rostrocaudal location on *Rxfp3* co-expression (ZI *Th*+: χ^2^_(3)_ = 15.48, *p* = .0014; LH *Th*+: χ^2^_(3)_ = 16.82, *p* < .001; LH *Pvalb*+: χ^2^_(3)_ = 503.72, *p* < .001; Figure 5F, J, K); however, none of the *post hoc* pairwise comparisons were statistically significant (all *p*s > .05). For ZI *Pvalb*+ cells, there was also a significant overall effect of rostrocaudal location on *Rxfp3* co-expression (χ^2^_(3)_ = 16.02, *p* = .0011). *Post hoc* pairwise comparisons revealed that the proportion of *Rxfp3*+/*Pvalb*+ cells was higher in the rZID/ZIV than in all other groups (ZIR, *p* = .011; iZID/ZIV, *p* = .002; ZIC, *p* < .001; Figure 5G). Notably, although the proportion of *Rxfp3*+ cells expressing *Pvalb* in the ZIC was ∼80%, the average proportion of *Pvalb*+ cells expressing *Rxfp3*+ in the ZIC was only ∼29%, suggesting that despite the high propensity of *Rxfp3*+/*Pvalb*+ cells in the area, *Rxfp3*+ expression does not sufficiently define *Pvalb* expression.

We then examined the spatial distribution of *Rxfp3*+/*Th*+, *Rxfp3*+/*Pvalb*+, and *Rxfp3*+/*Th*+/*Pvalb*+ cells across the LH and ZI. In the ZI, *Rxfp3*+/*Th*+ cells primarily congregated in the A13 area and were sparsely scattered across the rest of the ZI (Figure 6A – J). Additionally, the A13 was located more medially than its demarcation in the Mouse Brain Atlas, consistent with a previous report in the mouse (Negishi et al., 2020). Therefore, the mapping of the A13 area in Figure 6B - D shows these exceeded borders. *Rxfp3*+/*Pvalb*+ cells predominantly populated the ZIV, though they were ubiquitously present throughout the entire ZI (Figure 6A - J). In the LH, most *Rxfp3*+/*Th*+ cells occupied the medial LH, with fewer in more lateral areas (Figure 6K - V). Additionally, a cluster of *Rxfp3*+/*Th*+ cells was found surrounding the lateral aspect of the fornix in the intermediate/caudal aLH (Figure 6N - P). *Rxfp3*+/*Pvalb*+ cells were sparsely scattered throughout the LH, with no obvious patterns of expression (Figure 6K - V).

**Figure 6.**
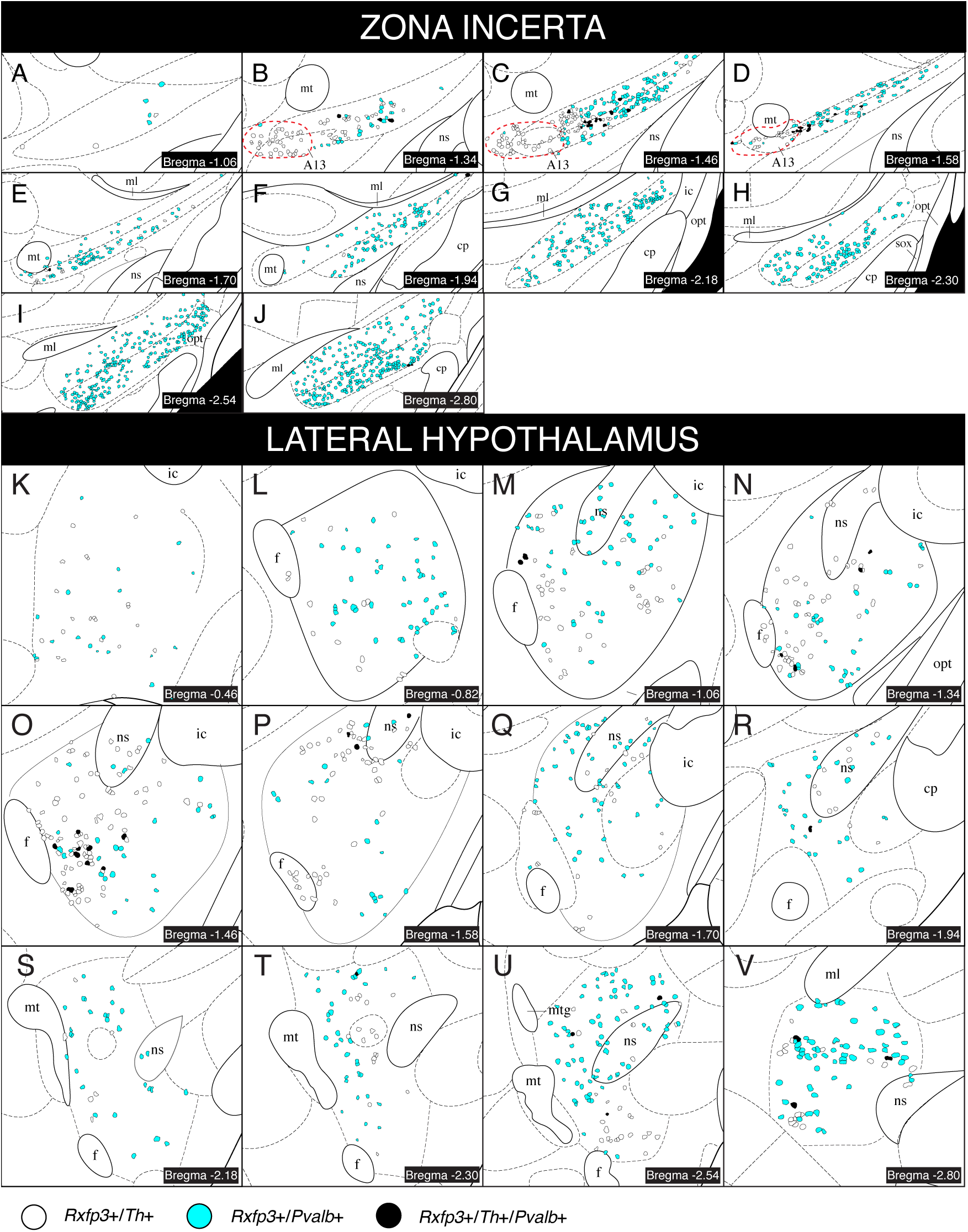
Schematics of *Rxfp3* co-expression with *Th* and *Pvalb* throughout the LH and ZI. *Rxfp3*+ cells that co-expressed *Th* (white dots), *Pvalb* (cyan dots), or both *Th* and *Pvalb* (black dots) were mapped onto plates within the ZI (A – J) and LH (K – V) from the Mouse Brain Atlas in Stereotaxic Coordinates (Paxinos & Franklin, 2001) from a representative RXFP3-Cre mouse. Note that mapping in B – D was performed beyond the published ZI borders due to the location of the A13 group (red dotted circles). Abbreviations: cp, cerebral peduncle; ic, internal capsule; ml, medial lemniscus; mt, mammillothalamic tract; mtg, mammillotegmental tract; ns, nigrostriatal bundle; opt, optic tract; sox, supraoptic decussation.

#### 3.2.3. Experiment 3: Somatostatin

ZI somatostatin-expressing (SST+) cells are sensitive to visual looming stimuli (Lin et al., 2023) and activating these cells is anxiogenic (Z. Li et al., 2021). Chemogenetic activation of LH SST+ cells increases general locomotor behaviour (Mickelsen et al., 2019). Consequently, we sought to determine whether LH/ZI^RXFP3^ cells express somatostatin by examining the colocalisation of *Rxfp3* with *Sst* mRNA. In the ZI, 14.6% (± 0.5%) of *Rxfp3*+ cells were *Sst*+ (Figure 7A, I), and 14.0% (± 2.1%) of *Sst*+ cells were *Rxfp3*+ (Figure 7F). In the LH, 14.2% (± 0.2%) of *Rxfp3*+ cells were *Sst*+ (Figure 7C), and 15.6% (± 1.6%) of *Sst*+ cells were *Rxfp3*+ (Figure 7H). Taken together, although there was evidence of some coexpression, RXFP3+ and SST+ populations in the LH and ZI are largely separate and only partially overlap.

**Figure 7.**
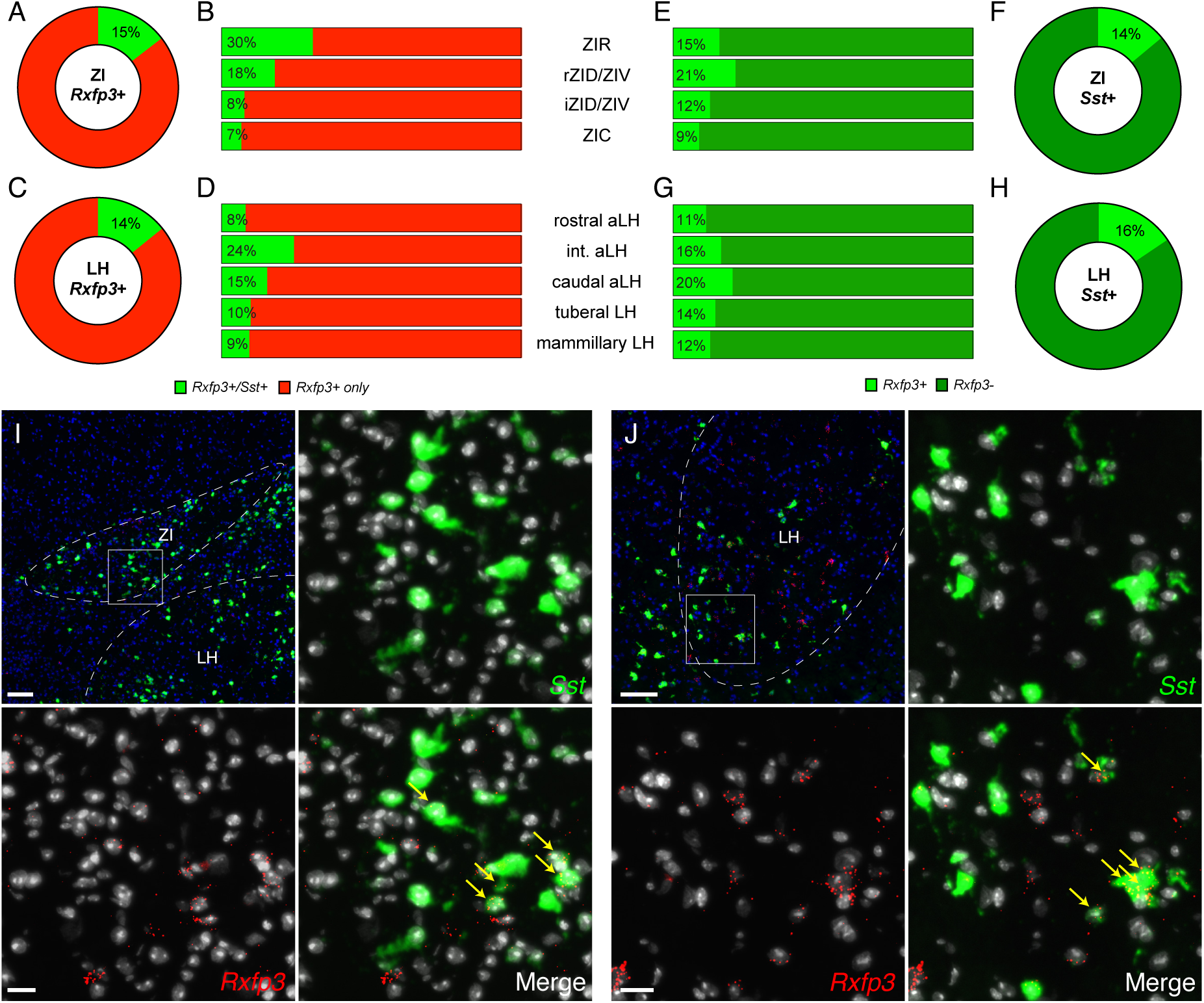
Co-localisation of *Rxfp3* with *Sst* throughout the ZI and LH. **(A, C)** Donut chart quantification of the mean proportion of *Rxfp3*+ cells expressing *Sst* (*Rxfp3*+/*Sst*+, light green), or not (*Rxfp3*+ only, red) in the ZI (A) and LH (C). **(B, D)** Distribution plots showing the mean proportion of *Rxfp3*+ cells expressing *Sst* or not but split across rostrocaudal location for the ZI (B) and LH (D). Reported means are adjusted values based on multinomial logistic regression. **(F, H)** Donut chart quantification of the mean proportion of *Sst*+ cells expressing *Rxfp3* (*Rxfp3*+, light green) or not (*Rxfp3*-, dark green) for both the ZI (F) and LH (H). **(E, G)** Distribution plots showing the mean proportion of *Sst*+ cells expressing *Rxfp3* or not but split across rostrocaudal location for the ZI (E) and the LH (G). Reported means are adjusted values based on binomial logistic regression. **(I, J)** Representative fluorescent photomicrograph of cells expressing *Rxfp3* and *Sst* mRNA relative to DAPI-stained nuclei in the ZI (I) and LH (J). Zoomed in images show the inset box in the top left panels. *Rxfp3*+/*Sst*+ cells are indicated by yellow arrows in the merge image. Main image scale bars = 100 µm. Inset image scale bars = 20 µm. All *n*s = 4 mice.

In the ZI, a binomial logistic regression revealed a significant overall effect of rostrocaudal location on *Rxfp3* composition (χ^2^_(3)_ = 145.8, *p* < .001). *Post hoc* multiple comparisons indicated that the proportion of *Rxfp3*+/*Sst*+ cells was significantly higher in the ZIR than in both the iZID/ZIV and ZIC (*p* < .001), and higher in the rZID/ZIV than in the ZIC (*p* < .001; Figure 7B). In the LH, a binomial logistic regression also revealed a significant overall effect of rostrocaudal location on *Rxfp3* composition (χ^2^_(3)_ = 52.71, *p* < .001). *Post hoc* multiple comparisons indicated that the proportion of *Rxfp3*+/*Sst*+ cells was significantly higher in the intermediate aLH compared to all other groups (all ps < .001), and higher in the caudal aLH compared to the rostral aLH, tuberal LH (ps < .001), and the mammillary LH (*p* = .013; Figure 7D). Additionally, we used binomial logistic regressions to test whether the proportion of *Sst*+ cells expressing *Rxfp3* changed along the rostrocaudal axis of the LH and ZI. For ZI *Sst*+ cells, there was a significant overall effect of rostrocaudal location on *Rxfp3*+ coexpression (χ^2^_(3)_ = 32.12, *p* < .001). *Post hoc* multiple comparisons indicated that the proportion of *Sst*+/*Rxfp3*+ cells was significantly higher in the ZIR and rZID/ZIV compared to the ZIC (ZIR vs ZIC, *p* = .045; rZID/ZIV vs ZIC, *p* = .003; Figure 7E). For LH *Sst*+ cells, there was a significant overall effect of rostrocaudal location on *Rxfp3*+ coexpression (χ^2^_(3)_ = 1270.53, *p* < .001). *Post hoc* multiple comparisons indicated that the proportion of *Sst*+/*Rxfp3*+ cells was significantly higher in the caudal aLH compared to the rostral aLH (*p* < .001) and mammillary LH (*p* = .002), higher in the intermediate aLH compared to the mammillary LH (*p* < .001), and higher in the tuberal LH compared with the rostral aLH (*p* = .001; Figure 7G).

We then examined the spatial distribution of *Rxfp3*+/*Sst*+ across the LH and ZI (Figure 8). In the ZIR, *Rxfp3*+/*Sst*+ cells were broadly distributed throughout the structure, with small clusters in the caudal A13 (Figure 8E, F). Expression was sparse throughout the rest of the ZI and did not cluster in any specific regions (Figure 8G – L). In the LH, some clusters of *Rxfp3*+/*Sst*+ cells were present in the intermediate aLH (Figure 8O, P), but no other distinct clusters were observed elsewhere in the LH.

**Figure 8.**
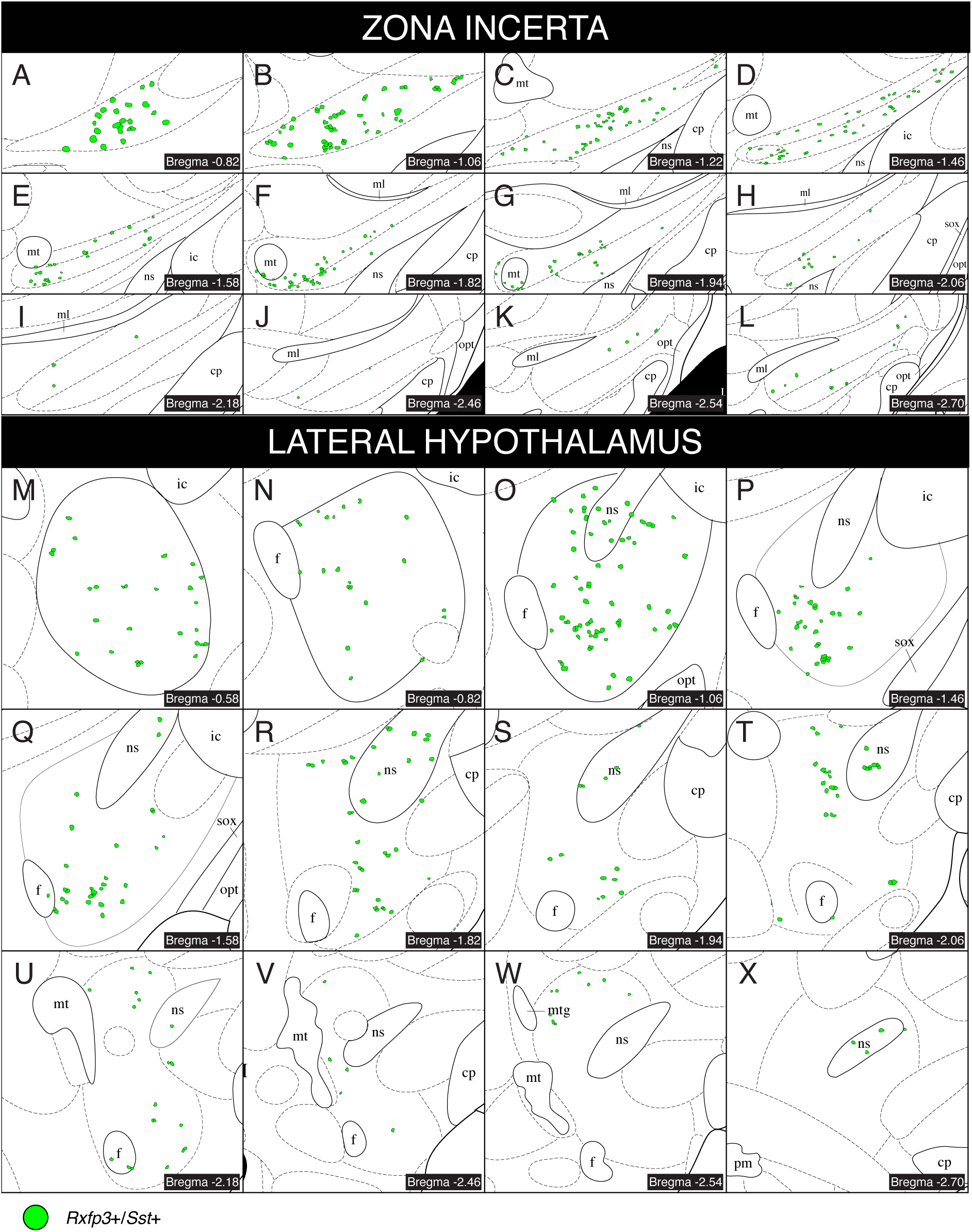
Schematics of *Rxfp3* co-expression with *Sst* throughout the LH and ZI. *Rxfp3*+ cells that co-expressed *Sst* (green dots), were mapped onto plates within the ZI (A – L) and LH (M – X) from the Mouse Brain Atlas in Stereotaxic Coordinates (Paxinos & Franklin, 2001) from a representative RXFP3-Cre mouse. Abbreviations: cp, cerebral peduncle; ic, internal capsule; ml, medial lemniscus; mt, mammillothalamic tract; mtg, mammillotegmental tract; ns, nigrostriatal bundle; opt, optic tract; sox, supraoptic decussation.

## 4. Discussion

Neuropeptides and their receptors have diverse modulatory roles across the central nervous system. Therefore, understanding their neuroanatomical properties is critical to fully appreciate their functional complexity. Here, we performed RNAscope *in situ* hybridisation to determine the spatial expression pattern and neurochemical phenotype of *Rxfp3*-expressing cells throughout the mouse LH and ZI. We demonstrated that *Rxfp3* is expressed across the rostrocaudal extent of both the LH and ZI and accounts for approximately one sixth of all DAPI-labelled nuclei in each structure. Neurochemical phenotyping of *Rxfp3*+ cells with *Gad1*, *Slc17a6*, *Pvalb*, *Th*, and *Sst* showed that LH/ZI *Rxfp3*+ cells co-express each marker to varying extents, generally proportional to their overall abundance within each structure (Figure 9). Overall, we have demonstrated that LH/ZI^RXFP3^ cells are neurochemically diverse, which may reflect their functional heterogeneity.

**Figure 9.**
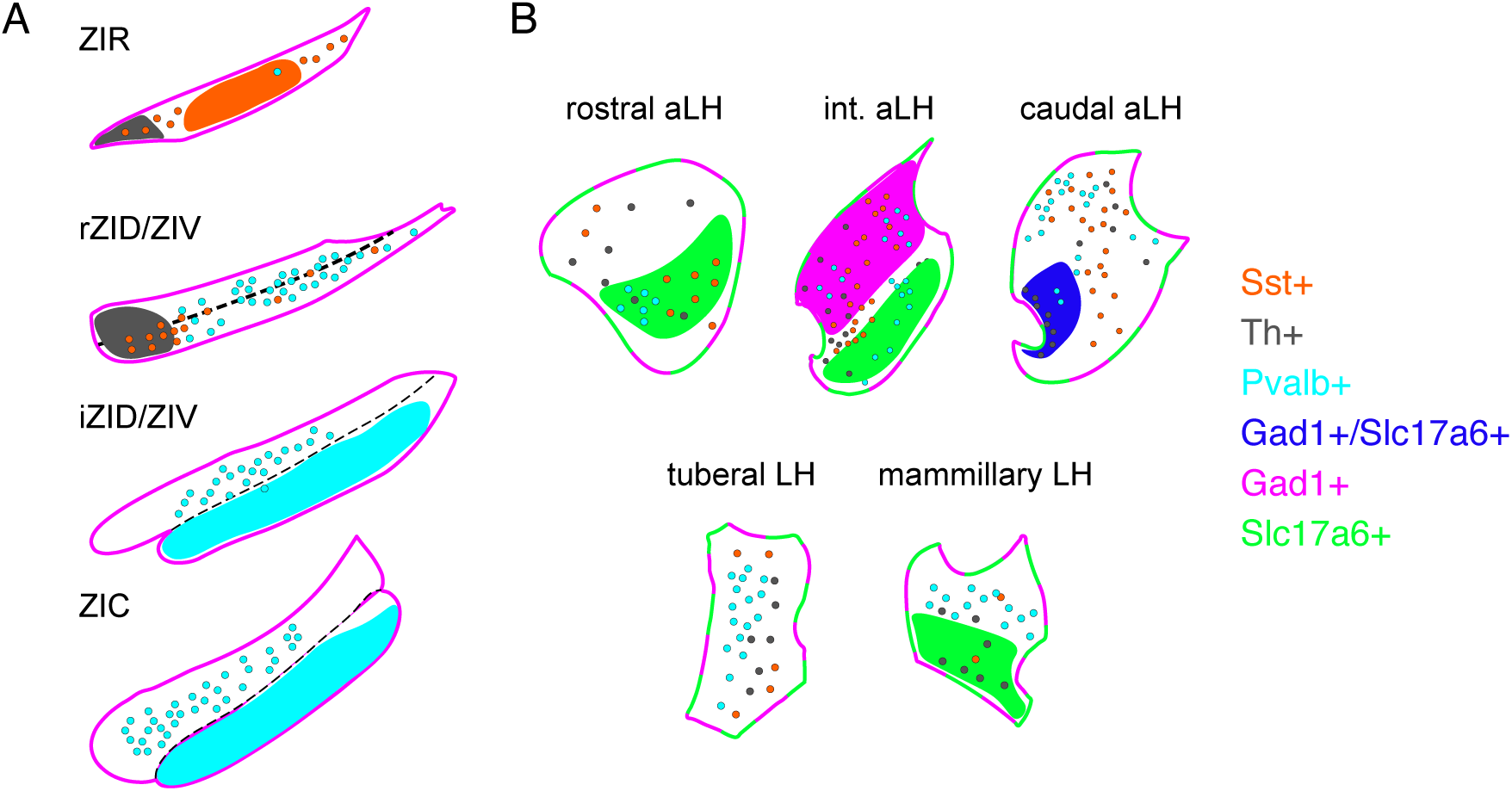
Summary of the *Rxfp3*+ co-expression profile across the LH/ZI. Schematic illustration of results for *Rxfp3*+ co-expression with *Sst*+ (orange), *Th*+ (grey), *Pvalb*+ (cyan), *Gad1*+ (pink), and *Slc17a6*+ (green) across the ZI (A) and LH (B). The pink outlines across the ZI rostrocaudal regions indicate that the majority of *Rxfp3*+ cells colocalised with *Gad1*+, while the green and pink outlines of the LH rostrocaudal regions indicate that *Rxfp3*+ cells colocalised with large proportions of *Gad1*+ and *Slc17a6*+. Coloured highlighted regions indicate area of high colocalisation density with the corresponding marker. Dots indicate relative densities of colocalisation in that area.

*Rxfp3* was widely expressed across the rostrocaudal extent of the LH and ZI. Prominent *Rxfp3*+ clusters were observed in the medial ZI surrounding the mammillothalamic tract, lateral ZIV, and in the caudal aLH surrounding the fornix, consistent with Allen Brain Cell Atlas data (Yao et al., 2023). However, a previous study reported that *Rxfp3* mRNA was scarcely present in the mouse ZI (Smith et al., 2010), and Allen Brain *in situ* hybridisation data show a much lower density of *Rxfp3* in the LH and ZI than observed presently (Lein et al., 2007). These discrepancies may reflect differences in *in situ* hybridisation techniques used across studies. Both RNAscope^™^ and multiplexed error-robust fluorescent *in situ* hybridisation (MERFISH; used in Yao et al., 2023) are highly sensitive techniques capable of detecting single mRNA molecules (Chen et al., 2015; F. Wang et al., 2012), making them suitable for detecting the low copy number of *Rxfp3* transcript observed in most LH/ZI cells. Traditional digoxigenin- and radioisotopic *in situ* hybridisation (used in Lein et al., 2007 and Smith et al., 2010, respectively) lack this sensitivity and thus may not have been able to resolve *Rxfp3* transcripts in the ZI (Gaspar & Ephrussi, 2015; F. Wang et al., 2012). A more recent study that examined the distribution of RXFP3+ cells using an RXFP3-Cre/YFP reporter line reported a moderate-to-high density of YFP+ cells in the ZI (Eraslan et al., 2024), which is more consistent with the current findings. They also noted that many regions with high YFP immunofluorescence showed no *Rxfp3* mRNA expression in Smith et al. 2010. Therefore, RNAscope™ or other more sensitive *in situ* hybridisation techniques, such as MERFISH, could be used to validate the presence of *Rxfp3* mRNA in these areas, given our findings.

Our data show that LH/ZI^RXFP3^ cells overlap with several neurochemically defined populations involved in various facets of fear learning and defensive behaviour. PV+ ZI neurons are necessary for the acquisition and storage of conditioned fear memories (Zhou et al., 2018), and their activation enhances active escape behaviour (X. Wang, Chou, Peng, et al., 2019). Optogenetic activation of TH+ A13 neurons projecting to the nucleus reuniens enhances fear extinction recall (Venkataraman et al., 2021). Optogenetic activation of SST+ ZI cells enhances defensive freezing to an innately threatening stimulus (Lin et al., 2023) and reduces time spent in the open arm of an elevated O-maze, indicating that their activity is anxiogenic (Z. Li et al., 2021). In the LH, optogenetic activation of glutamatergic projections to the lateral habenula is aversive (Lazaridis et al., 2019; Stamatakis et al., 2016), and GABAergic LH neurons are recruited during Pavlovian fear conditioning if animals have prior reward experience (Sharpe et al., 2021). Given that we recently demonstrated that chemogenetically activating LH/ZI^RXFP3^ cells following Pavlovian fear conditioning produced multiple fear-related behavioural phenotypes across animals (Richards et al., 2025), it is likely that we activated different proportions of these neurochemically defined subsets. Future research should aim to understand how these subpopulations of LH/ZI^RXFP3^ cells may influence various aspects of fear and defensive behaviours.

Our findings also support the view that classifying ZI neurons solely by GABAergic neurotransmission may hinder a clear understanding of ZI function. Studies that have functionally interrogated GABAergic ZI cells as a homogeneous population may have inadvertently captured multiple functionally distinct populations. Parsing the roles of GABAergic subtypes defined by their unique genetic or neurochemical signatures may be a more appropriate way forward to delineate ZI function. The utility of this approach is well demonstrated by recent work that has disentangled neurochemically defined GABAergic microcircuits in the central amygdala (CeA) that control the choice between flight and freezing in response to threats. Both corticotropin-releasing factor (CRF) expressing and SST-expressing CeA neurons are GABAergic, but CRF+ neurons are required to produce flight, while SST+ neurons are required to initiate freezing – two mutually exclusive behaviours (Fadok et al., 2017; Moscarello & Penzo, 2022; Tovote et al., 2016).

Our data also indicated that LH/ZI^RXFP3^ cells may overlap with other functional subpopulations. Though the functional literature surrounding neurochemically defined ZI subtypes remains scarce, ZI parvalbumin-expressing neurons have been implicated in pain (H. Wang et al., 2020), itch (J. Li et al., 2022), and grooming (Ge et al., 2024), while dopaminergic A13 neurons are linked to motor actions (Bisht et al., 2025; Garau et al., 2023), likely via their dense projections to the mesencephalic locomotor region (Bono et al., 2025; Sharma et al., 2018). The glutamatergic and GABAergic phenotype of many LH^RXFP3^ cells suggest that they may be involved in canonical LH functions, such as feeding and reward (Bonnavion et al., 2016). Further insight into the roles of LH^RXFP3^ cells will likely be gleaned by examining the distinct projection patterns of glutamatergic and GABAergic LH neurons, as well as their molecular profiles at a more granular level. For example, a recent single-cell transcriptomic study demonstrated that mouse LH neurons could be categorised into 15 distinct glutamatergic and 15 distinct GABAergic subtypes (Mickelsen et al., 2019), while spatiomolecular profiling of 24 genes in the LH showed that the region could be further subdivided into 9 subregions based on differential expression of discriminatory genes in glutamatergic and GABAergic subclasses (Y. Wang et al., 2021). Understanding which glutamatergic and GABAergic LH subtypes express RXFP3 is crucial to further elucidate its functions.

In experiments 1 and 2, a small proportion of *Rxfp3+*/*Slc17a6+*/*Gad1+* and *Rxfp3+*/*Th+*/*Pvalb+* cells were observed, respectively. To the best of our knowledge, *Th+*/*Pvalb+* cells in the LH and ZI have not been reported; their occurrence is possibly due to false-positive detection in areas of high marker overlap. However, one cluster of LH *Rxfp3*+/*Slc17a6*+/*Gad1*+ cells stands out, comprising 8% of the caudal aLH *Rxfp3*+ population, which is unlikely attributable to false-positive detection. This population may overlap with a vGlut2/VGAT co-expressing LH cluster recently identified by Wang and colleagues (2021). These findings raise the possibility of glutamate/GABA co-release from these cells, challenging the canonical understanding of excitatory versus inhibitory neurotransmission. Although the functional significance of these neurons is uncertain, the discrete clustering of these cells in the caudal aLH may imply an important regulatory function that is worth further interrogation.

A consistent finding across experiments was that *Rxfp3*+ expressing cells were a large subset of each neurochemical marker of interest. Furthermore, the proportion of *Rxfp3*+ cells expressing a particular marker of interest was commensurate with the marker’s relative abundance in that area. In other words, *Rxfp3* expression was generally unselective. ∼82% of ZI *Rxfp3*+ cells were *Gad1+,* and ∼2% were *Slc17a6*+, but the ZI is primarily a GABAergic nucleus (Benson et al., 1992; Kolmac & Mitrofanis, 1999), and *Gad1*+ and *Slc17a6*+ cells expressed *Rxfp3* in similar proportions. Although ∼80% of ZIC *Rxfp3*+ cells were *Pvalb*+ and only ∼7% of ZIR *Rxfp3*+ cells were *Pvalb*+, ZI PV+ cells primarily populate the caudal half of the ZI and are less abundant in the ZIR (Kolmac & Mitrofanis, 1999), and both ZIC and ZIR *Pvalb*+ were ∼28-29% *Rxfp3*+. ∼30% of ZIR *Rxfp3*+ cells were *Sst*+, a significantly higher proportion than in all other ZI sectors; however, ZI SST+ cells are primarily located within the ZIR (Lin et al., 2023), and ZIR *Sst*+ cells were *Rxfp3*+ in a similar proportion to all other sectors. Taken together, differences in the overall neurochemical composition of *Rxfp3*+ cells across the LH/ZI can be explained by the relative abundance of neurochemicals in the area rather than *by Rxfp3*+ cells showing selectivity for a particular neurochemical subclass. These findings are not unprecedented, as RXFP3+ cells in other brain regions have demonstrated similar properties. For example, the posterior bed nucleus of the stria terminalis contains a large population of RXFP3+ cells that express excitatory and inhibitory markers in relatively equal proportions (Ch’ng et al., 2019), consistent with the overall neurochemical composition of this area (Poulin et al., 2009). Similarly, Haidar and colleagues (2019) demonstrated that the most *Rxfp3*+ cells in the medial septum and diagonal band of Broca co-expressed vGAT, with only a small percentage expressing vGlut2, aligning with the overall abundance of vGAT and vGlut2 in these regions (Yao et al., 2023). Although *Rxfp3* mRNA is expressed in many other brain regions (Smith et al., 2010) whose neurochemical identities remain unexplored, our findings add to the consistent pattern that, across many brain regions, RXFP3 is not selective for any particular neurochemical subpopulation. This intriguing property may reflect the overall function of relaxin-3/RXFP3 signalling. The relaxin-3/RXFP3 neuropeptidergic system is described as an ascending arousal system with properties similar to the monoaminergic arousal systems (Smith et al., 2014). This is because, much like these systems, relaxin-3 is locally synthesised in a restricted brainstem area (the nucleus incertus; Burazin et al., 2002) and exhibits widespread ascending projections to regulate global behavioural states (Smith et al., 2014). Given that both the LH and ZI are considered global integrative hubs implicated in diverse functions that require high vigilance states (X. Wang, Chou, Zhang, et al., 2019), it is feasible for RXFP3 to be expressed on multiple neurochemically defined subtypes in the LH/ZI if its job is to regulate overall CNS arousal levels.

It should be noted that the current study examined male mice only. Although the literature is limited regarding sex-specific expression of neurochemical markers in the LH and ZI, sex-specific differences in relaxin-3 and RXFP3 expression have been observed elsewhere, such as in the nucleus incertus and amygdala (Meadows & Byrnes, 2015). Furthermore, relaxin-3/RXFP3 signalling has been shown to influence food intake and body weight in a sexually dimorphic manner (Calvez et al., 2015, 2017; Lenglos et al., 2013, 2014, 2015). Given the known role of the LH in regulating food intake and body weight, it is possible that RXFP3 expression is sexually dimorphic in this region, which could be relevant for future studies investigating the relaxin-3/RXFP3 system in this context.

## Contributions

BKR and CJP designed the experiment. BKR performed all experiments and wrote the manuscript. JHK, AJL, CJP acquired funds for the research. All authors reviewed and edited the manuscript.

## Data availability statement

The data that support the findings of this study are available from the corresponding author upon reasonable request.

## Supporting information

Abbreviations

## Acknowledgements

We thank the Central Animal Facility staff at Macquarie University for animal husbandry.

## References

Albert-Gascó, H., Ma, S., Ros-Bernal, F., Sánchez-Pérez, A. M., Gundlach, A. L., & Olucha-Bordonau, F. E. (2018). GABAergic neurons in the rat medial septal complex express relaxin-3 receptor (RXFP3) mRNA. Frontiers in Neuroanatomy, 11, 133–148. 10.3389/fnana.2017.00133

Bankhead, P., Loughrey, M. B., Fernández, J. A., Dombrowski, Y., McArt, D. G., Dunne, P. D., McQuaid, S., Gray, R. T., Murray, L. J., Coleman, H. G., James, J. A., Salto-Tellez, M., & Hamilton, P. W. (2017). QuPath: Open source software for digital pathology image analysis. Scientific Reports, 7(1), 16878. 10.1038/s41598-017-17204-5

Barbano, M. F., Wang, H.-L., Zhang, S., Miranda-Barrientos, J., Estrin, D. J., Figueroa-González, A., Liu, B., Barker, D. J., & Morales, M. (2020). VTA glutamatergic neurons mediate innate defensive behaviors. Neuron, 107(2), 368–382.e8. 10.1016/j.neuron.2020.04.024

Bathgate, R. A. D., Ivell, R., Sherwood, O. D., & Summers, R. J. (2006). International Union of Pharmacology LVII: Recommendations for the Nomenclature of Receptors for Relaxin Family Peptides. Pharmacological Reviews, 58(1), 7–31. 10.1124/pr.58.1.9

Bathgate, R. A. D., Samuel, C. S., Burazin, T. C. D., Layfield, S., Claasz, A. A., Reytomas, I. G. T., Dawson, N. F., Zhao, C., Bond, C., Summers, R. J., Parry, L. J., Wade, J. D., & Tregear, G. W. (2002). Human Relaxin Gene 3 (*H3*) and the Equivalent Mouse Relaxin (*M3*) Gene: NOVEL MEMBERS OF THE RELAXIN PEPTIDE FAMILY. Journal of Biological Chemistry, 277(2), 1148–1157. 10.1074/jbc.M107882200

Benson, D. L., Isackson, P. J., Gall, C. M., & Jones, E. G. (1992). Contrasting patterns in the localization of glutamic acid decarboxylase and Ca2+ /calmodulin protein kinase gene expression in the rat central nervous system. Neuroscience, 46(4), 825–849. 10.1016/0306-4522(92)90188-8

Bisht, A., Badenhorst, C., Kiss, Z. H. T., Murari, K., & Whelan, P. J. (2025). Deep brain stimulation of A13 region evokes robust locomotory response in rats. Journal of Neurophysiology, 133(5), 1594–1606. 10.1152/jn.00019.2025

Bonnavion, P., Mickelsen, L. E., Fujita, A., de Lecea, L., & Jackson, A. C. (2016). Hubs and spokes of the lateral hypothalamus: Cell types, circuits and behaviour: LHA cell types and circuits. The Journal of Physiology, 594(22), 6443–6462. 10.1113/JP271946

Bono, B. S., Negishi, K., Dumiaty, Y., Ponce-Ruiz, M. S., Akinbode, T. C., Baker, K. S., Spencer, C. D. P., Mejia, E., Guirguis, M., Hebert, A. J., Khan, A. M., & Chee, M. J. (2025). Brainwide projections of mouse dopaminergic zona incerta neurons. Journal of Comparative Neurology, 533(3), e70039. 10.1002/cne.70039

Braga, A., Chiacchiaretta, M., Pellerin, L., Kong, D., & Haydon, P. G. (2024). Astrocytic metabolic control of orexinergic activity in the lateral hypothalamus regulates sleep and wake architecture. Nature Communications, 15(1), 5979. 10.1038/s41467-024-50166-7

Burazin, T. C. D., Bathgate, R. A. D., Macris, M., Layfield, S., Gundlach, A. L., & Tregear, G. W. (2002). Restricted, but abundant, expression of the novel rat gene-3 (R3) relaxin in the dorsal tegmental region of brain. Journal of Neurochemistry, 82(6), 1553–1557. 10.1046/j.1471-4159.2002.01114.x

Calvez, J., de Ávila, C., & Timofeeva, E. (2017). Sex-specific effects of relaxin-3 on food intake and body weight gain: Sex-specific effects of relaxin-3. British Journal of Pharmacology, 174(10), 1049–1060. 10.1111/bph.13530

Calvez, J., Lenglos, C., de Ávila, C., Guèvremont, G., & Timofeeva, E. (2015). Differential effects of central administration of relaxin-3 on food intake and hypothalamic neuropeptides in male and female rats: Sex-specific effects of relaxin-3. *Genes*, Brain and Behavior, 14(7), 550–563. 10.1111/gbb.12236

Calvigioni, D., Fuzik, J., Le Merre, P., Slashcheva, M., Jung, F., Ortiz, C., Lentini, A., Csillag, V., Graziano, M., Nikolakopoulou, I., Weglage, M., Lazaridis, I., Kim, H., Lenzi, I., Park, H., Reinius, B., Carlén, M., & Meletis, K. (2023). Esr1+ hypothalamic-habenula neurons shape aversive states. Nature Neuroscience, 26(7), 1245–1255. 10.1038/s41593-023-01367-8

Cezario, A. F., Ribeiro-Barbosa, E. R., Baldo, M. V. C., & Canteras, N. S. (2008). Hypothalamic sites responding to predator threats—The role of the dorsal premammillary nucleus in unconditioned and conditioned antipredatory defensive behavior. European Journal of Neuroscience, 28(5), 1003–1015. 10.1111/j.1460-9568.2008.06392.x

Chen, K. H., Boettiger, A. N., Moffitt, J. R., Wang, S., & Zhuang, X. (2015). Spatially resolved, highly multiplexed RNA profiling in single cells. Science, 348(6233), aaa6090. 10.1126/science.aaa6090

Ch’ng, S. S., Fu, J., Brown, R. M., Smith, C. M., Hossain, M. A., McDougall, S. J., & Lawrence, A. J. (2019). Characterization of the relaxin family peptide receptor 3 system in the mouse bed nucleus of the stria terminalis. Journal of Comparative Neurology, 527(16), 2615–2633. 10.1002/cne.24695

Chou, X., Wang, X., Zhang, Z., Shen, L., Zingg, B., Huang, J., Zhong, W., Mesik, L., Zhang, L. I., & Tao, H. W. (2018). Inhibitory gain modulation of defense behaviors by zona incerta. Nature Communications, 9(1), 1151. 10.1038/s41467-018-03581-6

Contesse, T., Bektash, B. Y., Graziano, M., Forastieri, C., Contestabile, A., Hahne, S., Jung, F., Nikolakopoulou, I., Rubino, E., Cao, X., Skara, V., Mantas, I., Giatrellis, S., Carlén, M., Sandberg, R., Calvigioni, D., & Meletis, K. (2025). A striosomal accumbens pathway drives stereotyped behavior through an aversive Esr1+ hypothalamic-habenula circuit. Science Advances, 11(47), 1–17. 10.1126/sciadv.adx9450

De Sousa, A. F., Cowansage, K. K., Zutshi, I., Cardozo, L. M., Yoo, E. J., Leutgeb, S., & Mayford, M. (2019). Optogenetic reactivation of memory ensembles in the retrosplenial cortex induces systems consolidation. Proceedings of the National Academy of Sciences, 116(17), 8576–8581. 10.1073/pnas.1818432116

Devarakonda, K., & Kenny, P. J. (2017). Energy balance: Lateral hypothalamus hoards food memories. Current Biology, 27(16), R803–R805. 10.1016/j.cub.2017.06.082

Dielenberg, R. A., Hunt, G. E., & McGregor, I. S. (2001). ‘When a rat smells a cat’: The distribution of Fos immunoreactivity in rat brain following exposure to a predatory odor. Neuroscience, 104(4), 1085–1097. 10.1016/S0306-4522(01)00150-6

Eraslan, I. M., Egberts-Brugman, M., Read, J. L., Voglsanger, L. M., Samarasinghe, R. M., Hamilton, L., Dhar, P., Williams, R. J., Walker, L. C., Ch’ng, S., Lawrence, A. J., Walker, A. J., Dean, O. M., Gundlach, A. L., & Smith, C. M. (2024). Neuroanatomical distribution of fluorophores within adult RXFP3 cre-tdTomato/YFP mouse brain. Biochemical Pharmacology, 225, 116265. 10.1016/j.bcp.2024.116265

Fadok, J. P., Krabbe, S., Markovic, M., Courtin, J., Xu, C., Massi, L., Botta, P., Bylund, K., Müller, C., Kovacevic, A., Tovote, P., & Lüthi, A. (2017). A competitive inhibitory circuit for selection of active and passive fear responses. Nature, 542(7639), 96–100. 10.1038/nature21047

Garau, C., Hayes, J., Chiacchierini, G., McCutcheon, J. E., & Apergis-Schoute, J. (2023). Involvement of A13 dopaminergic neurons in prehensile movements but not reward in the rat. Current Biology, 33(22), 4786–4797.e4. 10.1016/j.cub.2023.09.044

Gaspar, I., & Ephrussi, A. (2015). Strength in numbers: Quantitative single-molecule RNA detection assays. WIREs Developmental Biology, 4(2), 135–150. 10.1002/wdev.170

Ge, J., Ren, P., Tian, B., Li, J., Qi, C., Huang, Q., Ren, K., Hu, E., Mao, H., Zang, Y., Wu, S., Xue, Q., & Wang, W. (2024). Ventral zona incerta parvalbumin neurons modulate sensory-induced and stress-induced self-grooming via input-dependent mechanisms in mice. iScience, 27(7), 110165. 10.1016/j.isci.2024.110165

Gross, C. T., & Canteras, N. S. (2012). The many paths to fear. Nature Reviews Neuroscience, 13(9), 651–658. 10.1038/nrn3301

Hahn, J. D., Sporns, O., Watts, A. G., & Swanson, L. W. (2019). Macroscale intrinsic network architecture of the hypothalamus. Proceedings of the National Academy of Sciences, 116(16), 8018–8027. 10.1073/pnas.1819448116

Haidar, M., Tin, K., Zhang, C., Nategh, M., Covita, J., Wykes, A. D., Rogers, J., & Gundlach, A. L. (2019). Septal GABA and Glutamate Neurons Express RXFP3 mRNA and Depletion of Septal RXFP3 Impaired Spatial Search Strategy and Long-Term Reference Memory in Adult Mice. Frontiers in Neuroanatomy, 13, 30. 10.3389/fnana.2019.00030

Ho, P., Hsiao, F., Chiu, S., Lee, S., & Yau, H. (2023). A nigroincertal projection mediates aversion and enhances coping responses to potential threat. The FASEB Journal, 37(12), e23322. 10.1096/fj.202201989RR

Kania, A., Gugula, A., Grabowiecka, A., de Ávila, C., Blasiak, T., Rajfur, Z., Lewandowski, M. H., Hess, G., Timofeeva, E., Gundlach, A. L., & Blasiak, A. (2017). Inhibition of oxytocin and vasopressin neuron activity in rat hypothalamic paraventricular nucleus by relaxin-3-RXFP3 signalling: Relaxin-3-RXFP3 modulation of rat paraventricular nucleus activity. The Journal of Physiology, 595(11), 3425–3447. 10.1113/JP273787

Kim, D. J., Lee, A. S., Yttredahl, A. A., Gómez-Rodríguez, R., & Anderson, B. J. (2017). Repeated threat (without direct harm) alters metabolic capacity in select regions that drive defensive behavior. Neuroscience, 353, 106–118. 10.1016/j.neuroscience.2017.04.012

Kolmac, C., & Mitrofanis, J. (1999). Distribution of various neurochemicals within the zona incerta: An immunocytochemical and histochemical study. Anatomy and Embryology, 199(3), 265–280. 10.1007/s004290050227

Laing, B. T., Anderson, M. S., Bonaventura, J., Jayan, A., Sarsfield, S., Gajendiran, A., Michaelides, M., & Aponte, Y. (2023). Anterior hypothalamic parvalbumin neurons are glutamatergic and promote escape behavior. Current Biology, 33(15), 3215–3228.e7. 10.1016/j.cub.2023.06.070

Lazaridis, I., Tzortzi, O., Weglage, M., Märtin, A., Xuan, Y., Parent, M., Johansson, Y., Fuzik, J., Fürth, D., Fenno, L. E., Ramakrishnan, C., Silberberg, G., Deisseroth, K., Carlén, M., & Meletis, K. (2019). A hypothalamus-habenula circuit controls aversion. Molecular Psychiatry, 24(9), 1351–1368. 10.1038/s41380-019-0369-5

Lecca, S., Meye, F. J., Trusel, M., Tchenio, A., Harris, J., Schwarz, M. K., Burdakov, D., Georges, F., & Mameli, M. (2017). Aversive stimuli drive hypothalamus-to-habenula excitation to promote escape behavior. eLife, 6, e30697. 10.7554/eLife.30697

Lein, E. S., Hawrylycz, M. J., Ao, N., Ayres, M., Bensinger, A., Bernard, A., Boe, A. F., Boguski, M. S., Brockway, K. S., Byrnes, E. J., Chen, L., Chen, L., Chen, T.-M., Chi Chin, M., Chong, J., Crook, B. E., Czaplinska, A., Dang, C. N., Datta, S., … Jones, A. R. (2007). Genome-wide atlas of gene expression in the adult mouse brain. Nature, 445(7124), 168–176. 10.1038/nature05453

Lenglos, C., Calvez, J., & Timofeeva, E. (2015). Sex-Specific Effects of Relaxin-3 on Food Intake and Brain Expression of Corticotropin-Releasing Factor in Rats. Endocrinology, 156(2), 523–533. 10.1210/en.2014-1743

Lenglos, C., Mitra, A., Guèvremont, G., & Timofeeva, E. (2013). Sex differences in the effects of chronic stress and food restriction on body weight gain and brain expression of CRF and relaxin-3 in rats: Sex difference in body weight regulation. *Genes*, Brain and Behavior, 12(4), 370–387. 10.1111/gbb.12028

Lenglos, C., Mitra, A., Guèvremont, G., & Timofeeva, E. (2014). Regulation of expression of relaxin-3 and its receptor RXFP3 in the brain of diet-induced obese rats. Neuropeptides, 48(3), 119–132. 10.1016/j.npep.2014.02.002

Li, J., Bai, Y., Liang, Y., Zhang, Y., Zhao, Q., Ge, J., Li, D., Zhu, Y., Cai, G., Tao, H., Wu, S., & Huang, J. (2022). Parvalbumin Neurons in Zona Incerta Regulate Itch in Mice. Frontiers in Molecular Neuroscience, 15, 843754. 10.3389/fnmol.2022.843754

Li, Z., Rizzi, G., & Tan, K. R. (2021). Zona incerta subpopulations differentially encode and modulate anxiety. Science Advances, 7(37), eabf6709. 10.1126/sciadv.abf6709

Lin, S., Zhu, M.-Y., Tang, M.-Y., Wang, M., Yu, X.-D., Zhu, Y., Xie, S.-Z., Yang, D., Chen, J., & Li, X.-M. (2023). Somatostatin-Positive Neurons in the Rostral Zona Incerta Modulate Innate Fear-Induced Defensive Response in Mice. Neuroscience Bulletin, 39(2), 245–260. 10.1007/s12264-022-00958-y

Liu, C., Eriste, E., Sutton, S., Chen, J., Roland, B., Kuei, C., Farmer, N., Jörnvall, H., Sillard, R., & Lovenberg, T. W. (2003). Identification of Relaxin-3/INSL7 as an Endogenous Ligand for the Orphan G-protein-coupled Receptor GPCR135. Journal of Biological Chemistry, 278(50), 50754–50764. 10.1074/jbc.M308995200

Ma, S., Bonaventure, P., Ferraro, T., Shen, P.-J., Burazin, T. C. D., Bathgate, R. A. D., Liu, C., Tregear, G. W., Sutton, S. W., & Gundlach, A. L. (2007). Relaxin-3 in GABA projection neurons of nucleus incertus suggests widespread influence on forebrain circuits via G-protein-coupled receptor-135 in the rat. Neuroscience, 144(1), 165–190. 10.1016/j.neuroscience.2006.08.072

Meadows, K. L., & Byrnes, E. M. (2015). Sex- and age-specific differences in relaxin family peptide receptor expression within the hippocampus and amygdala in rats. Neuroscience, 284, 337–348. 10.1016/j.neuroscience.2014.10.006

Mendes-Gomes, J., Motta, S. C., Passoni Bindi, R., De Oliveira, A. R., Ullah, F., Baldo, M. V. C., Coimbra, N. C., Canteras, N. S., & Blanchard, D. C. (2020). Defensive behaviors and brain regional activation changes in rats confronting a snake. Behavioural Brain Research, 381, 112469. 10.1016/j.bbr.2020.112469

Mickelsen, L. E., Bolisetty, M., Chimileski, B. R., Fujita, A., Beltrami, E. J., Costanzo, J. T., Naparstek, J. R., Robson, P., & Jackson, A. C. (2019). Single-cell transcriptomic analysis of the lateral hypothalamic area reveals molecularly distinct populations of inhibitory and excitatory neurons. Nature Neuroscience, 22(4), 642–656. 10.1038/s41593-019-0349-8

Mitrofanis, J. (2005). Some certainty for the “zone of uncertainty”? Exploring the function of the zona incerta. Neuroscience, 130(1), 1–15. 10.1016/j.neuroscience.2004.08.017

Moeyaert, B., Holt, G., Madangopal, R., Perez-Alvarez, A., Fearey, B. C., Trojanowski, N. F., Ledderose, J., Zolnik, T. A., Das, A., Patel, D., Brown, T. A., Sachdev, R. N. S., Eickholt, B. J., Larkum, M. E., Turrigiano, G. G., Dana, H., Gee, C. E., Oertner, T. G., Hope, B. T., & Schreiter, E. R. (2018). Improved methods for marking active neuron populations. Nature Communications, 9(1), 4440. 10.1038/s41467-018-06935-2

Moscarello, J. M., & Penzo, M. A. (2022). The central nucleus of the amygdala and the construction of defensive modes across the threat-imminence continuum. Nature Neuroscience, 25(8), 999–1008. 10.1038/s41593-022-01130-5

Navarro-Sánchez, M., Gil-Miravet, I., Montero-Caballero, D., Bathgate, R. A. D., Hossain, M. A., Castillo-Gómez, E., Gundlach, A. L., & Olucha-Bordonau, F. E. (2024). Modulation of contextual fear acquisition and extinction by acute and chronic relaxin-3 receptor (RXFP3) activation in the rat retrosplenial cortex. Biochemical Pharmacology, 225, 116264. 10.1016/j.bcp.2024.116264

Negishi, K., Payant, M. A., Schumacker, K. S., Wittmann, G., Butler, R. M., Lechan, R. M., Steinbusch, H. W. M., Khan, A. M., & Chee, M. J. (2020). Distributions of hypothalamic neuron populations coexpressing tyrosine hydroxylase and the vesicular GABA transporter in the mouse. Journal of Comparative Neurology, 528(11), 1833–1855. 10.1002/cne.24857

Paxinos, G., & Franklin, K. B. J. (2001). The Mouse Brain in Stereotaxic Coordinates (2nd ed.). Academic Press.

Petzold, A., Van Den Munkhof, H. E., Figge-Schlensok, R., & Korotkova, T. (2023). Complementary lateral hypothalamic populations resist hunger pressure to balance nutritional and social needs. Cell Metabolism, 35(3), 456–471.e6. 10.1016/j.cmet.2023.02.008

Poulin, J.-F., Arbour, D., Laforest, S., & Drolet, G. (2009). Neuroanatomical characterization of endogenous opioids in the bed nucleus of the stria terminalis. Progress in Neuro-Psychopharmacology and Biological Psychiatry, 33(8), 1356–1365. 10.1016/j.pnpbp.2009.06.021

Richards, B. K., Ch’ng, S. S., Simon, A. B., Pang, T. Y., Kim, J. H., Lawrence, A. J., & Perry, C. J. (2025). Relaxin family peptide receptor 3 (RXFP3) expressing cells in the zona incerta/lateral hypothalamus augment behavioural arousal. Journal of Neurochemistry, 169(1), e16217. 10.1111/jnc.16217

Richards, B. K., Kilby, A. I. J., Cornish, J. L., Kim, J. H., Lawrence, A. J., & Perry, C. J. (2026). Efferent projections of topographically distinct relaxin family peptide receptor-3 (RXFP3) lateral hypothalamus/zona incerta cells. bioRxiv. 10.64898/2026.01.15.699820

Rytova, V., Ganella, D. E., Hawkes, D., Bathgate, R. A. D., Ma, S., & Gundlach, A. L. (2019). Chronic activation of the relaxin-3 receptor on GABA neurons in rat ventral hippocampus promotes anxiety and social avoidance. Hippocampus, 29(10), 905–920. 10.1002/hipo.23089

Schroeder, A., Pardi, M. B., Keijser, J., Dalmay, T., Groisman, A. I., Schuman, E. M., Sprekeler, H., & Letzkus, J. J. (2023). Inhibitory top-down projections from zona incerta mediate neocortical memory. Neuron, 111(5), 727–738.e8. 10.1016/j.neuron.2022.12.010

Sharma, S., Kim, L. H., Mayr, K. A., Elliott, D. A., & Whelan, P. J. (2018). Parallel descending dopaminergic connectivity of A13 cells to the brainstem locomotor centers. Scientific Reports, 8(1), 7972. 10.1038/s41598-018-25908-5

Sharpe, M. J., Batchelor, H. M., Mueller, L. E., Gardner, M. P. H., & Schoenbaum, G. (2021). Past experience shapes the neural circuits recruited for future learning. Nature Neuroscience, 24(3), 391–400. 10.1038/s41593-020-00791-4

Siemian, J. N., Borja, C. B., Sarsfield, S., Kisner, A., & Aponte, Y. (2019). Lateral hypothalamic fast-spiking parvalbumin neurons modulate nociception through connections in the periaqueductal gray area. Scientific Reports, 9(1), 12026. 10.1038/s41598-019-48537-y

Smith, C. M., Shen, P.-J., Banerjee, A., Bonaventure, P., Ma, S., Bathgate, R. A. D., Sutton, S. W., & Gundlach, A. L. (2010). Distribution of relaxin-3 and RXFP3 within arousal, stress, affective, and cognitive circuits of mouse brain. The Journal of Comparative Neurology, 518(19), 4016–4045. 10.1002/cne.22442

Smith, C. M., Walker, A. W., Hosken, I. T., Chua, B. E., Zhang, C., Haidar, M., & Gundlach, A. L. (2014). Relaxin-3/RXFP3 networks: An emerging target for the treatment of depression and other neuropsychiatric diseases? Frontiers in Pharmacology, 5. 10.3389/fphar.2014.00046

Soya, S., Takahashi, T. M., McHugh, T. J., Maejima, T., Herlitze, S., Abe, M., Sakimura, K., & Sakurai, T. (2017). Orexin modulates behavioral fear expression through the locus coeruleus. Nature Communications, 8(1), 1606. 10.1038/s41467-017-01782-z

Stamatakis, A. M., Van Swieten, M., Basiri, M. L., Blair, G. A., Kantak, P., & Stuber, G. D. (2016). Lateral Hypothalamic Area Glutamatergic Neurons and Their Projections to the Lateral Habenula Regulate Feeding and Reward. The Journal of Neuroscience, 36(2), 302–311. 10.1523/JNEUROSCI.1202-15.2016

Swanson, L. (1987). The hypothalamus. In A. Björklund, T. Hökfelt, & L. Swanson (Eds.), Integrated Systems of the CNS. Part I (Vol. 5). Elsevier.

Swanson, L. W. (2000). Cerebral hemisphere regulation of motivated behavior. Brain Research, 0–52. 10.1016/S0006-8993(00)02905-X

Teegala, S. B., Sarkar, P., Siegel, D. M., Sheng, Z., Hao, L., Bello, N. T., De Lecea, L., Beck, K. D., & Routh, V. H. (2023). Lateral hypothalamus hypocretin/orexin glucose-inhibited neurons promote food seeking after calorie restriction. Molecular Metabolism, 76, 101788. 10.1016/j.molmet.2023.101788

Tovote, P., Esposito, M. S., Botta, P., Chaudun, F., Fadok, J. P., Markovic, M., Wolff, S. B. E., Ramakrishnan, C., Fenno, L., Deisseroth, K., Herry, C., Arber, S., & Lüthi, A. (2016). Midbrain circuits for defensive behaviour. Nature, 534(7606), 206–212. 10.1038/nature17996

Venkataraman, A., Brody, N., Reddi, P., Guo, J., Gordon Rainnie, D., & Dias, B. G. (2019). Modulation of fear generalization by the zona incerta. Proceedings of the National Academy of Sciences, 116(18), 9072–9077. 10.1073/pnas.1820541116

Venkataraman, A., & Dias, B. G. (2023). Expanding the canon: An inclusive neurobiology of thalamic and subthalamic fear circuits. Neuropharmacology, 226, 109380. 10.1016/j.neuropharm.2022.109380

Venkataraman, A., Hunter, S. C., Dhinojwala, M., Ghebrezadik, D., Guo, J., Inoue, K., Young, L. J., & Dias, B. G. (2021). Incerto-thalamic modulation of fear via GABA and dopamine. Neuropsychopharmacology, 46(9), 1658–1668. 10.1038/s41386-021-01006-5

Viden, A., Ch’ng, S. S., Walker, L. C., Shesham, A., Hamilton, S. M., Smith, C. M., & Lawrence, A. J. (2022). Organisation of enkephalin inputs and outputs of the central nucleus of the amygdala in mice. Journal of Chemical Neuroanatomy, 125, 102167. 10.1016/j.jchemneu.2022.102167

Voglsanger, L. M., Read, J., Ch’ng, S. S., Zhang, C., Eraslan, I. M., Gray, L., Rivera, L. R., Hamilton, L. D., Williams, R., Gundlach, A. L., & Smith, C. M. (2021). Differential Level of RXFP3 Expression in Dopaminergic Neurons Within the Arcuate Nucleus, Dorsomedial Hypothalamus and Ventral Tegmental Area of RXFP3-Cre/tdTomato Mice. Frontiers in Neuroscience, 14, 594818. 10.3389/fnins.2020.594818

Walker, L. C., Hand, L. J., Letherby, B., Huckstep, K. L., Campbell, E. J., & Lawrence, A. J. (2021). Cocaine and amphetamine regulated transcript (CART) signalling in the central nucleus of the amygdala modulates stress-induced alcohol seeking. Neuropsychopharmacology, 46(2), 325–333. 10.1038/s41386-020-00807-4

Wang, F., Flanagan, J., Su, N., Wang, L.-C., Bui, S., Nielson, A., Wu, X., Vo, H.-T., Ma, X.-J., & Luo, Y. (2012). RNAscope: A Novel in Situ RNA Analysis Platform for Formalin-Fixed, Paraffin-Embedded Tissues. The Journal of Molecular Diagnostics, 14(1), 22–29. 10.1016/j.jmoldx.2011.08.002

Wang, H., Dong, P., He, C., Feng, X.-Y., Huang, Y., Yang, W.-W., Gao, H.-J., Shen, X.-F., Lin, S., Cao, S.-X., Lian, H., Chen, J., Yan, M., & Li, X.-M. (2020). Incerta-thalamic Circuit Controls Nocifensive Behavior via Cannabinoid Type 1 Receptors. Neuron, 107(3), 538–551.e7. 10.1016/j.neuron.2020.04.027

Wang, X., Chou, X., Peng, B., Shen, L., Huang, J. J., Zhang, L. I., & Tao, H. W. (2019). A cross-modality enhancement of defensive flight via parvalbumin neurons in zonal incerta. eLife, 8, e42728. 10.7554/eLife.42728

Wang, X., Chou, X., Zhang, L. I., & Tao, H. W. (2019). Zona Incerta: An Integrative Node for Global Behavioral Modulation. Trends in Neurosciences, 43(2), 82–87. 10.1016/j.tins.2019.11.007

Wang, Y., Eddison, M., Fleishman, G., Weigert, M., Xu, S., Wang, T., Rokicki, K., Goina, C., Henry, F. E., Lemire, A. L., Schmidt, U., Yang, H., Svoboda, K., Myers, E. W., Saalfeld, S., Korff, W., Sternson, S. M., & Tillberg, P. W. (2021). EASI-FISH for thick tissue defines lateral hypothalamus spatio-molecular organization. Cell, 184(26), 6361–6377.e24. 10.1016/j.cell.2021.11.024

Wu, K., Wang, D., Wang, Y., Tang, P., Li, X., Pan, Y., Tao, H. W., Zhang, L. I., & Liang, F. (2023). Distinct circuits in anterior cingulate cortex encode safety assessment and mediate flexibility of fear reactions. Neuron, 111(22), S0896627323006190. 10.1016/j.neuron.2023.08.008

Yao, Z., Van Velthoven, C. T. J., Kunst, M., Zhang, M., McMillen, D., Lee, C., Jung, W., Goldy, J., Abdelhak, A., Aitken, M., Baker, K., Baker, P., Barkan, E., Bertagnolli, D., Bhandiwad, A., Bielstein, C., Bishwakarma, P., Campos, J., Carey, D., … Zeng, H. (2023). A high-resolution transcriptomic and spatial atlas of cell types in the whole mouse brain. Nature, 624(7991), 317–332. 10.1038/s41586-023-06812-z

Zeng, H., & Sanes, J. R. (2017). Neuronal cell-type classification: Challenges, opportunities and the path forward. Nature Reviews Neuroscience, 18(9), 530–546. 10.1038/nrn.2017.85

Zhang, W., Zhang, N., Sakurai, T., & Kuwaki, T. (2009). Orexin neurons in the hypothalamus mediate cardiorespiratory responses induced by disinhibition of the amygdala and bed nucleus of the stria terminalis. Brain Research, 1262, 25–37. 10.1016/j.brainres.2009.01.022

Zhou, M., Liu, Z., Melin, M. D., Ng, Y. H., Xu, W., & Südhof, T. C. (2018). A central amygdala to zona incerta projection is required for acquisition and remote recall of conditioned fear memory. Nature Neuroscience, 21(11), 1515–1519. 10.1038/s41593-018-0248-4

