## Supplementary material for "Neurochemical phenotype of relaxin family peptide receptor-3 (RXFP3) lateral hypothalamus/zona incerta cells": Abbreviations

**Table 1. List of abbreviations**

| anterior lateral hypothalamus | aLH |
| --- | --- |
| intermediate part of the dorsal/ventral zona incerta | iZID/ZIV |
| lateral hypothalamus | LH |
| multiplexed error-robust fluorescent in situ hybridisation | MERFISH |
| phosphate-buffered saline | PBS |
| paraformaldehyde | PFA |
| parvalbumin | PV |
| room temperature | RT |
| relaxin family peptide receptor 3 | RXFP3 |
| rostral part of the dorsal/ventral zona incerta | rZID/ZIV |
| somatostatin | SST |
| tyrosine hydroxylase | TH |
| zona incerta | ZI |
| caudal part of the dorsal/ventral zona incerta and caudal zona incerta | ZIC |
| dorsal zona incerta | ZID |
| rostral zona incerta | ZIR |
| ventral zona incerta | ZIV |
